# Exogenous chromosomes reveal how sequence composition drives chromatin assembly, activity, folding and compartmentalization

**DOI:** 10.1101/2022.12.21.520625

**Authors:** Christophe Chapard, Léa Meneu, Jacques Serizay, Alex Westbrook, Etienne Routhier, Myriam Ruault, Amaury Bignaud, Agnès Thierry, Géraldine Gourgues, Carole Lartigue, Aurèle Piazza, Angela Taddei, Frédéric Beckouët, Julien Mozziconacci, Romain Koszul

**Author notes:** These authors contributed equally. Univ Lyon, ENS, UCBL, CNRS, INSERM, Laboratory of Biology and Modelling of the Cell, UMR5239, U 1210, F-69364, Lyon, France.

## Abstract

Genomic sequences co-evolve with DNA-associated proteins to ensure the multiscale folding of long DNA molecules into functional chromosomes. In eukaryotes, different molecular complexes organize the chromosome’s hierarchical structure, ranging from nucleosomes and cohesin- mediated DNA loops to large scale chromatin compartments. To explore the relationships between the DNA sequence composition and the spontaneous loading and activity of these DNA-associated complexes in the absence of co-evolution, we characterized chromatin assembly and activity in yeast strains carrying exogenous bacterial chromosomes that diverged from eukaryotic sequences over 1.5 billion years ago. We show that nucleosome assembly, transcriptional activity, cohesin-mediated looping, and chromatin compartmentalization can occur in a bacterial chromosome with a largely divergent sequence integrated in a eukaryotic host, and that the chromatinization of bacterial chromosomes is highly correlated with their sequence composition. These results are a step forward in understanding how foreign sequences are interpreted by a host nuclear machinery during natural horizontal gene transfers, as well as in synthetic genomics projects.

## Introduction

Genome sequence composition, broadly defined by its GC%, polynucleotide frequencies, DNA motifs and repeats, varies widely between species as well as within individual genomes^1,2^. In eukaryotes, the sequence composition is known to correlate with: 1) chromatin composition, which includes nucleosome formation and binding of structural and functional proteins to DNA ^3^, 2) chromatin activity, such as transcription and replication ^4^, and 3) functional 3D organization of the genome into loops and compartments ^5,6^. For instance in mammals GC-rich regions are enriched in actively transcribed sequences and in chromatin loops mediated by the structural maintenance of chromosomes (SMC) cohesin, and genomic loci with different sequence composition tend to coalesce in distinct compartments ^7^. These relationships between sequence and chromatin composition activities and folding reflect their continuous coevolution over millions of years.

Disruptive variations in sequence composition can also emerge naturally during evolution, e.g. through transfer of genetic material across species by horizontal gene transfers or introgression, in viral infections ^8–11^, or even artificially, e.g. by introducing chromosome-long DNA molecules into chassis microbial strains or cell lines ^12–14^. Such transfer can lead to the long-term integration of foreign DNA whose sequence composition strongly diverges from their host’s genome (e.g. the introgression of a 12% GC-divergent 1Mb sequence in *Lachancea kluyveri* ^15^). Once integrated, these sequences are organized and processed by chromatin-associated proteins of the host genome, obeying new rules under which they have not coevolved. How a eukaryotic host can successfully package, fold and regulate the activity of chromosome-long exogenous bacterial DNA sequences, and the importance of the sequence composition, remain largely unknown. Addressing this question would improve our understanding of the evolutionary processes and sequence determinants involved in chromatin folding and activity.

Here we explored whether and how foreign bacterial full-length chromosomes acquire chromatin features when integrated in a eukaryotic nucleus, and the relationship between their sequence composition and their chromatinization. To this end, we investigated the behavior of two individual bacterial chromosomes with different sequence composition, artificially introduced into the *S. cerevisiae* genome ^16,17^. We profiled nucleosome assembly, polymerase and cohesin binding and activities, as well as the 3D chromosome organization during the cell cycle. We show that bacterial chromosomes with different sequence composition present different profiles of chromatin composition and activities, eventually leading to the spontaneous formation of active / inactive chromosomal compartments, similar to the ones observed in complex multicellular organisms. Using neural networks (NNs), we further show that eukaryotic sequence determinants are sufficient to predict the chromatin composition and activity of exogenous chromosomes integrated in the yeast genome. Consequently, this study suggests that the fate of any DNA molecule introduced into a given cellular context, including its organization from nucleosome positioning up to 3D folding and transcriptional activity, is governed by rules that are both deterministic and predictable.

## Results

### Adaptation of supernumerary bacteria chromosomes to yeast

To investigate large sequences which have not evolved in a eukaryotic context, we exploited *S. cerevisiae* strains carrying an extra 17^th^ circular chromosome made either from the *Mycoplasma mycoides* subsp. *mycoides* (referred to as “Mmyco”) or *Mycoplasma pneumoniae* (“Mpneumo”) genomes containing a yeast centromeric sequence and an autonomous replication sequence (ARS) ^18^ (**Methods**; **Table S1**). While the GC content of *S. cerevisiae* is 38%, the GC content of the Mmyco and Mpneumo chromosomes 24% (GC-poor) and 40% (GC-neutral), respectively (**Fig. 1a**). Dinucleotide composition, estimated by the dinucleotide odds ratio ρ*(XY) ^19^, also largely differs between yeast and bacterial chromosomes (**Extended Data** Fig. 1a).

**Figure 1.**
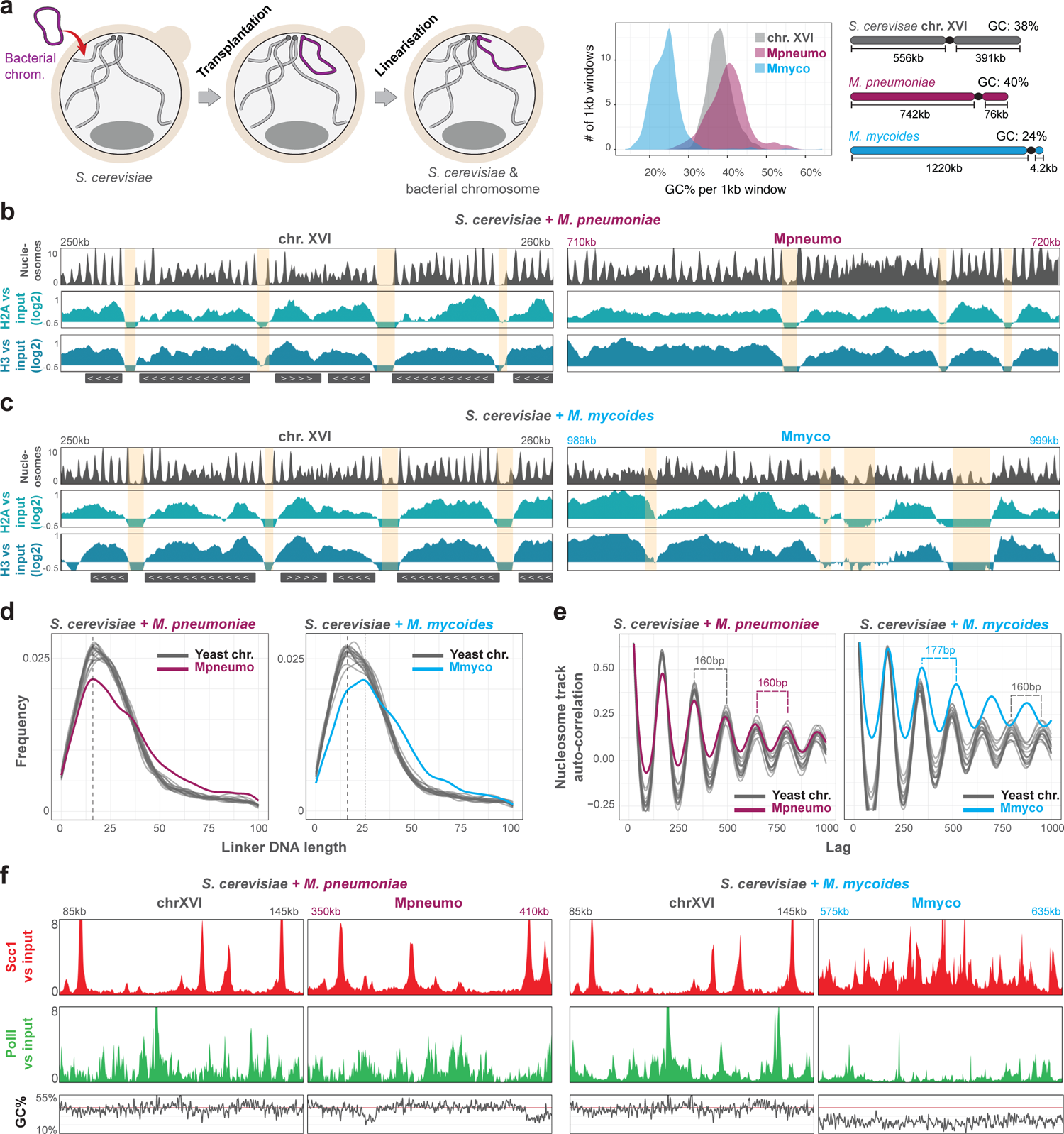
Chromatin composition of bacterial chromosomes in yeast. **a**, Schematic representation of the conversion from circular to linear chromosomes integrated in yeast. The purple and blue colors represent the *M. pneumoniae* (Mpneumo) and *M. mycoides* (Mmyco) bacterial sequence in all figures, respectively. Right: distribution of GC% for 1kb windows over yeast chromosome XVI, *M. pneumoniae* and *M. mycoides* chromosomes. **b**, Nucleosomal track (grey, see **Methods**) and H2A and H3 ChIP-seq (shades of blue, IP vs input, log2) profiles obtained in the Mpneumo strain (*S. cerevisiae + M. pneumoniae*). 10kb-long genomic windows from the chromosome XVI (left: 250kb-260kb) and the Mpneumo chromosome (right: 710kb-720kb) are shown at the same scale. Genomic loci highlighted in yellow correspond to nucleosome-depleted regions. **c**, Same as **b** in the Mmyco strain (*S. cerevisiae + M. mycoides*). 10kb-long genomic windows from chromosome XVI (left: 250kb-260kb) and the Mmyco chromosome (right: 989kb-999kb) are shown at the same scale. **d**, Frequency of nucleosome linker DNA length in Mpneumo and Mmyco strains. For each strain, the distribution is calculated for each chromosome separately. The dashed line indicates 14 bp and the dotted line indicates 25 bp. **e**, Auto-correlation of the nucleosome track in Mpneumo and Mmyco strains, computed for each chromosome separately (lag <= 1000). Nucleosome repeat length (NRL) is indicated for yeast, Mpneumo and Mmyco chromosomes. **f**, Scc1 (red) and RNA Pol II (green) ChIP-seq profiles (IP vs input, log2) obtained in the Mpneumo (left) or in the Mmyco (right) strains. For each strain, 60kb-long genomic windows from the bacterial chromosome and the chromosome XVI are shown, at the same scale. GC% in sliding 1kb windows is shown below the ChIP-seq profiles.

We linearized these chromosomes, added yeast telomeres at the extremities (**Fig. 1a; Extended Data** Fig. 1b; **Methods**), and investigated their replication and cohesion using split- dot assay and marker frequency analysis (**Extended Data** Fig. 1c,d and **Supplementary Results**; **Methods**). Overall, these chromosomes do not impose a significant fitness cost on their eukaryotic host, with each strain displaying similar growth rates compared to their parental strains in selective media (**Extended Data** Fig. 1e). They also have an estimated segregation rate similar to a centromeric plasmid ^20^ (**Extended Data** Fig. 1f).

### Spontaneous chromatinization of bacterial chromosomes in a eukaryotic context

We first assessed whether integrated Mmyco and Mpneumo chromosomes were able to chromatinize by performing MNase-seq and H3 and H2A ChIP-seq experiments (Methods; **Fig. 1b, c; Extended Data** Fig. 2a). Strikingly, both bacterial chromosomes were able to form nucleosomes and local nucleosome-depleted regions (NDR; **Fig. 1b, c; Extended Data** Fig. 2b-d). Nucleosomes over the Mpneumo chromosome have structural features comparable to yeast nucleosomes, with a linker DNA of ∼14 bp and a nucleosomal repeat length (NRL) of 160 bp (**Fig. 1d,e**). In contrast, nucleosomes in the Mmyco chromosome have a longer linker DNA and thus are more spaced than in yeast or Mpneumo chromosomes, with an NRL of 177 bp (**Fig. 1d,e)**. Two sequence features may explain this: first, poly(dA) and poly(dT) tracks, known to promote NDR formation, are over-represented along the Mmyco genome. Secondly, the AA/TT 10-bp periodicity, which facilitates DNA bending around histone cores, is reduced (**Extended Data** Fig. 2e,f). The analysis of MNase-seq time course experiments also revealed that Mmmyco nucleosomal fragments are more rapidly degraded by MNase than yeast or Mpneumo nucleosomal fragments (**Extended Data** Fig. 2c,d), reflecting either the “fragile” nature of AT-rich nucleosomes or the MNAse bias for these sequences ^21,22^.

We further profiled the chromatin composition of Mmyco and Mpneumo chromosomes by performing RNA polymerase II (PolII) and cohesin (Scc1) ChIP-seq experiments (**Fig. 1f, Extended Data Fig. 2g-j**). We found that the Scc1 and PolII binding profiles along the Mpneumo chromosome appear similar to wild-type (WT) yeast chromosomes, with discrete Scc1 peaks (**Fig. 1f, Extended Data** Fig. 2h,i) preferentially located in nucleosome-depleted and PolII-enriched regions (**Extended Data** Fig. 2j). In contrast, Scc1 and PolII binding profiles in the Mmyco strain show significant differences. On the one hand, Scc1 is strongly enriched over the whole Mmyco chromosome, but does not form the distinct peaks observed on endogenous chromosomes (**Fig. 1f, Extended Data** Fig. 2h,i). Interestingly, Scc1 levels also appear strongly reduced at centromeres of endogenous *S. cerevisiae* chromosomes (**Extended Data** Fig. 2g), suggesting that the Mmyco chromosome titrates cohesins enriched over yeast centromeres. On the other hand, the PolII profile is greatly reduced along the Mmyco chromosome, compared with yeast chromosomes.

These results show that large exogenous bacterial chromosomes placed in a eukaryotic context spontaneously adopt eukaryotic chromatin composition features: histones, Pol II and cohesins all bind bacterial DNA irrespectively of their sequence composition. However, striking differences in the chromatin profiles led us to define two chromatin types: (1) “Y” (yeast-like) chromatin landscape, found over Mpneumo (whose 40% GC content is close to the native *S. cerevisiae* GC content), and (2) “U” (for Unconventional) chromatin, found over Mmyco (with a low 24% GC content), featuring less packed nucleosomes, a reduced Pol II enrichment, and a broad binding of cohesins across the entire chromosome.

Finally, we find that a convolutional neural network trained on yeast chromosome sequences was able to predict nucleosome, Scc1 and PolII binding profiles along integrated bacterial chromosomes (**Supplementary Results** and **Extended Data** Fig. 3; **Methods**). This indicates that the complex, underlying eukaryotic sequence determinants are successfully reused to generate these different chromatin landscapes along integrated bacterial chromosomes.

### Transcriptional activity of bacterial genomes in a yeast context

We next explored Y and U chromatin activities by first performing total, stranded RNA-seq (**Fig. 2a,b; Method**). Consistent with PolII ChIP-seq profiles (**Fig. 1f**), we find that the Y chromatin type on Mpneumo is transcribed to levels similar to those of endogenous yeast chromosomes (**Fig. 2a**). However, Mpneumo transcription tracks are significantly longer than yeast genes (4.9kb vs 3.4kb, p-value < 2e-4, two-sided Student’s t-test) and do not systematically display clear boundaries (**Fig. 2b**). They also do not preferentially occur over bacterial gene bodies nor seem to initiate at bacterial promoters, also consistent with an absence of PolII enrichment at these loci (**Extended Data** Fig. 4a-c).

**Figure 2.**
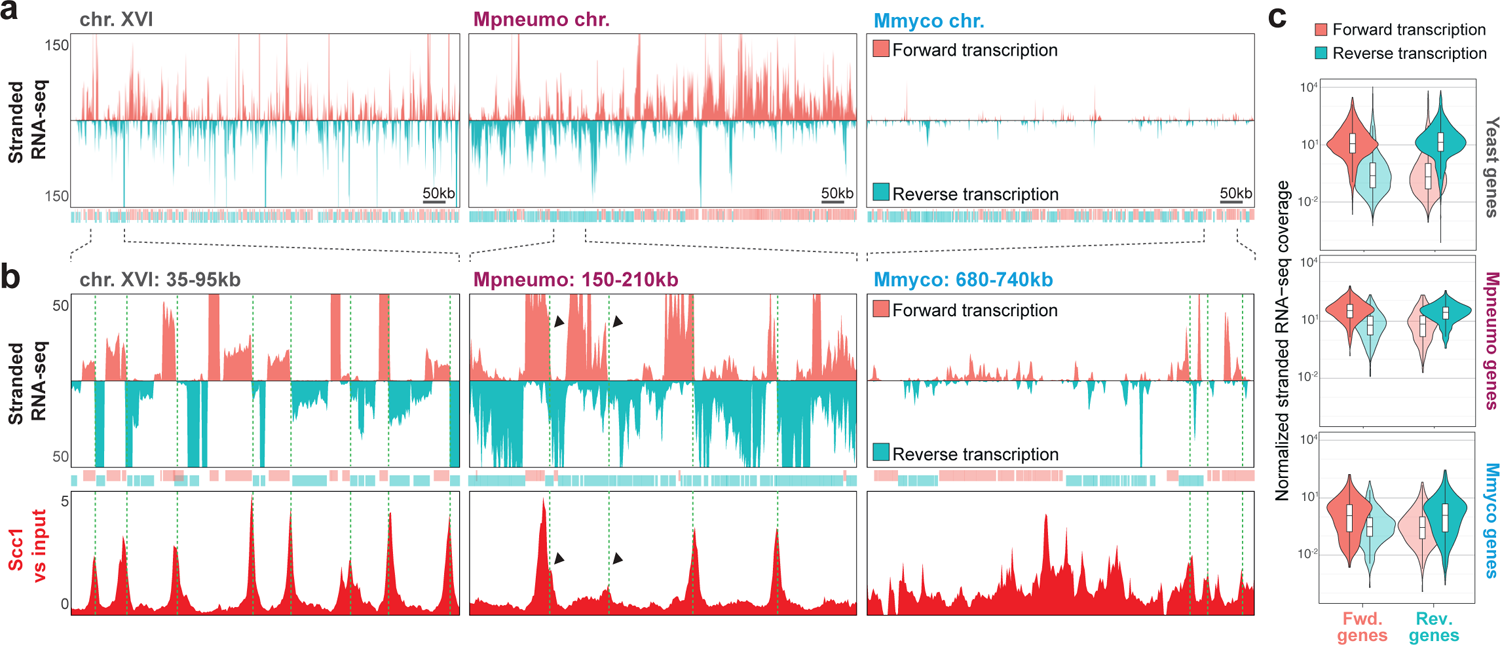
Expression of exogenic bacterial sequences in budding yeast. **a,** Stranded RNA-seq profiles along yeast chromosome XVI or along bacterial chromosomes Mpneumo and Mmyco in the corresponding strains. Forward (pink) and reverse (turquoise) genes along yeast or bacterial sequences are indicated as transparent segments under the tracks. Pink and turquoise represent forward and reverse transcription, respectively. **b,** Top: stranded RNA-seq profiles along a 60kb window along yeast chromosome XVI, Mpneumo or Mmyco. Bottom: Scc1 (cohesin) deposition profiles of the corresponding loci. Green dotted lines indicate identified loci of convergent transcription (see Methods). **c,** Forward and reverse RNA-seq coverage of forward- and reverse-oriented yeast or bacterial genes. Scores were normalized by the length of each genomic feature.

In sharp contrast, the Mmyco U chromatin type is only sparsely and lowly transcribed (**Fig. 2a,b**), again in good agreement with the reduced levels of PolII deposition (**Fig. 1f**). This transcriptional inactivity is not associated with the recruitment of the yeast silencing complex SIR Sir2/3/4 ^23^ (**Extended Data** Fig. 4d).

Intriguingly, the transcription tracks orientation along bacterial chromosomes follows, on average, the orientation of the bacterial genes annotated along these sequences, although the transcription machinery has diverged billions years ago (**Fig. 2c**, see the orientation of genes and stranded tracks in **Fig. 2a,b**). This correlates with the over-representation of A (G) compared to T (C) on the coding strand, both detected in yeast genes and bacterial genes (**Extended Data** Fig. 4e, see **Discussion**).

### Inactive U chromatin forms a compartment into the nuclear space in G1

To assess the 3D organization of Y and U chromatin types, we performed capture of chromosome conformation (Hi-C) ^24^ experiments on the Mmyco or Mpneumo strains arrested either in G1 or G2/M (**Fig. 3a, b**).

**Figure 3.**
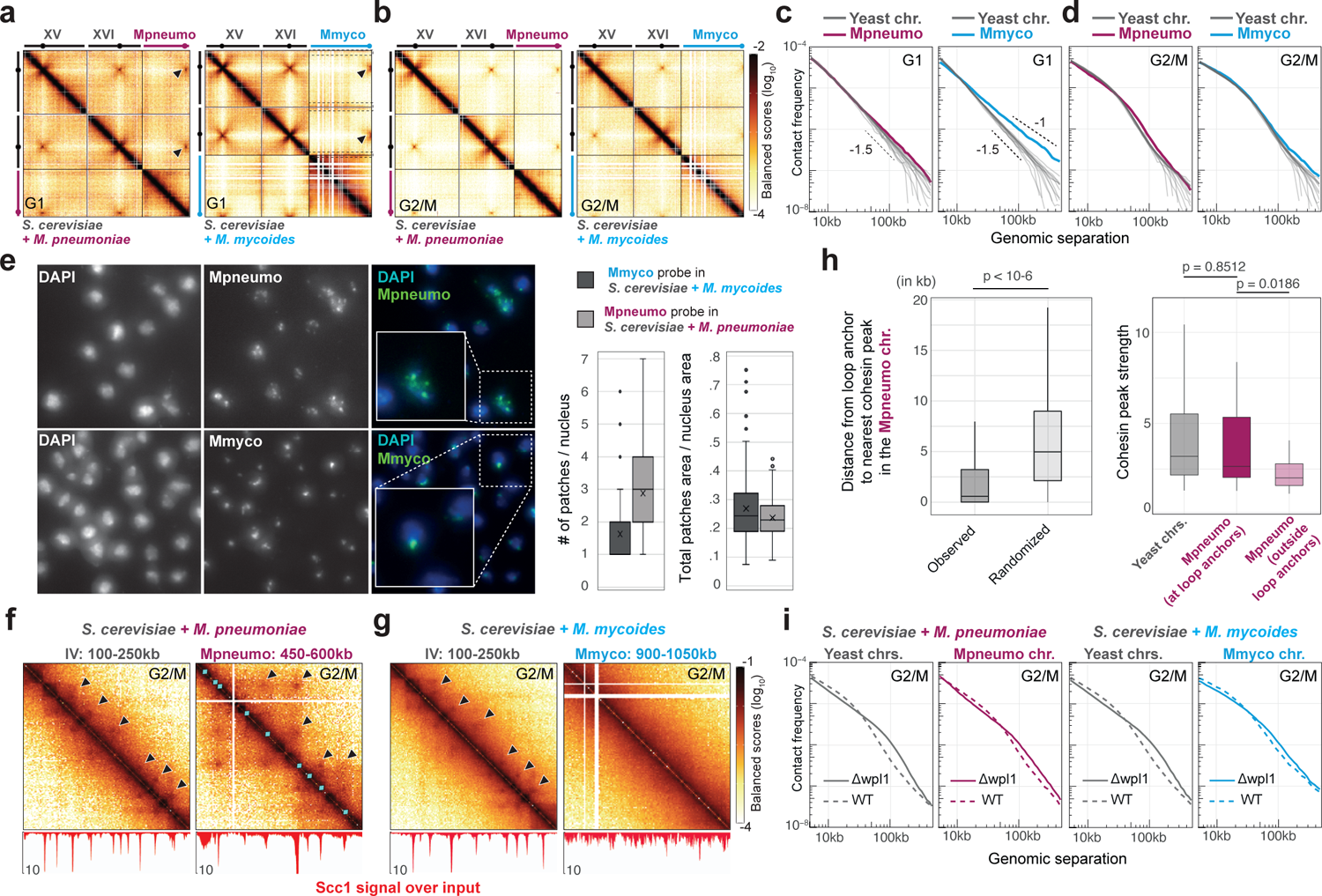
Folding of exogenic bacterial sequences within the yeast nucleus. **a, b,** Hi-C contact maps of representative endogenous and of Mmyco and Mpneumo bacterial chromosomes in G1 (**a**) and G2/M (**b**) (4kb resolution). **c**, **d**, Contact frequency (p) as a function of genomic distance (s) plots of endogenous yeast chromosomes (long arms) and of Mmyco and Mpneumo bacterial chromosomes in G1 (**c**) and G2/M (**d**). **e,** Left: FISH imaging. Representative field of either (top) Mpneumo or (bottom) Mmyco fixed cells labeled with DAPI (left panel) and hybridized with a fluorescent probe generated from either the Mpneumo or Mmyco chromosome, respectively. Right: For each probe, number of patches detected per nucleus and surface occupied by these patches relative to the whole nucleus surface (Methods). **f, g**, Top: magnification of 150kb windows from Hi-C contact maps in G2/M from either an endogenous or the bacterial chromosome in Mpneumo (**f**) and Mmyco (**g**) strains (1kb resolution). Bottom: Scc1 ChIP-seq deposition profile. Black arrowheads: loops. Cyan diamonds: Scc1 peaks positions. **h,** Left: distance between chromatin loop anchors and the nearest Scc1 peak in Mpneumo (with and without a random shuffle of peak positions). Right: Scc1 peak strengths in yeast or Mpneumo chromosome, near (< 1kb) or outside loop anchors (p-values from two-sided Student’s t-test). **i,** Distance-dependent contact frequency in endogenous yeast and in bacterial chromosomes (left, Mpneumo; right, Mmyco), in WT (dashed) and in Δwpl1 mutants.

In G1-arrested cells, yeast chromosomes exhibit the expected Rabl configuration, with clustered centromeres and subtelomeric regions, resulting in a dotted pattern in Hi-C contact maps ^25,26^. For both Mmyco and Mpneumo chromosomes, the added yeast centromeric sequence clusters with the native yeast centromeres. However, the two bacterial chromosomes exhibit very distinct structural characteristics (**Fig. 3a**). The Y-type Mpneumo chromosome behaves as an endogenous *S. cerevisiae* chromosome (**Fig. 3a**), with similar trans-contacts (**Extended Data** Fig. 5a) and long-range cis-contacts (**Fig. 3c**), and a slope of the averaged contact probability (p) as a function of the genomic distance (*s)* close to −1.5, a value corresponding to the typical polymer structure observed in simulations (**Fig. 3c**) ^27^. In addition, DNA-FISH labeling of Mpneumo DNA reveals an extended and contorted structure within the intranuclear space, confirming the intermixing of the chromosome with the actively transcribed native chromosomes (**Fig. 3e, Extended Data** Fig. 5c,d).

In contrast, in the U-type Mmyco chromosome, (1) only few contacts were detected with the endogenous chromosomes (**Extended Data** Fig. 5a), and predominantly with the 32 yeast subtelomeric regions (**Fig. 3a**, dotted rectangles; **Extended Data** Fig. 5b); (2) the slope of the p(s) curve also changes dramatically towards a value of −1 at shorter distances, corresponding to a polymer globule (**Fig. 3c**) ^24^; (3) DNA-FISH labeling of the Mmyco chromosome confirmed the globular conformation of the chromosome and revealed its preferential position at the nuclear periphery, reflecting its proximity with sub/telomeres (**Fig. 3e; Extended Data** Fig. 5c,d). These results show that inactive, U chromatin forms a compartment in the nuclear space segregated from the active endogenous chromosomal set.

### Cohesin compacts similarly both exogenous bacterial chromosomes during G2/M

We then generated Hi-C contact maps of both strains synchronized in G2/M. In yeast, chromosome compaction in G2/M-arrested cells reflect cohesin-mediated chromatin folding into arrays of loops, with cohesin peaks corresponding to loop anchors ^28,29^. This compaction results in a typical p(s) curve that presents an inflection point around the average loop length ^25^. The decrease of interchromosomal contacts (**Fig. 3b, Extended Data** Fig. 5a) and of p(s) (**Fig. 3d, Extended Data** Fig. 5e,f) observed for both native and bacterial chromosomes indicate a similar mitotic compaction of both Y and U chromatin types.

Yeast and Mpneumo chromosomes exhibit focal contact Scc1 enrichments corresponding to cohesin-anchored loops (black triangles on **Fig. 3f**) ^28^. We found that cohesin accumulates at sites of convergent transcription not only, as expected, on yeast chromosomes ^30,31^, but also on Mpneumo (**Fig. 2b**, green dotted lines, **Extended Data** Fig. 5g), with strong cohesin peaks associated with sharper transcriptional convergence (**Fig. 2b**, arrows; **Extended Data** Fig. 5h,i). Using Chromosight, we identified 59 loops across the Mpneumo chromosome, spanning over slightly longer genomic distances (28 kb on average) than the loops found along yeast sequences (22 kb) (**Extended Data** Fig. 5j; **Methods**), and with strong Scc1 peaks positioned close to anchors, with a median distance of 566 bp (**Fig. 3f, h**). The dotted grid pattern in Mpneumo is reminiscent of multiple DNA loops observed along mammalian interphase chromosomes ^32^, suggesting that some loop anchors have the ability to be involved in loops of various sizes, a phenomenon not observed across native *S. cerevisiae* chromosomes (**Fig. 3f**).

On the other hand, no visible discrete loops were observed along the Mmyco chromosome where cohesins cover inactive U chromatin regions without clear peaks, probably reflecting the absence of (convergent) transcription (**Fig. 3g**) (**Fig. 1f** and **Fig. 2b**). These results suggest that cohesin can bind DNA and promote compaction in absence of active transcription, but doest accumulate at discrete positions.

Cohesin-mediated loops along yeast chromosomes are formed through loop extrusion (LE), a process by which cohesin complexes organize DNA by capturing and gradually enlarging small loops ^33,34^. To test whether LE compacts bacterial genomes, we performed Hi-C in absence of Wpl1 (human Wapl), which impairs cohesin removal and results in longer loops readily visible by a shift of the inflection point in the p(s) curve ^28,34^. In absence of Wpl1, we observed increased long-range contacts in all chromosomes, including Mmyco and Mpneumo (**Fig. 3i; Extended Data** Fig. 5k), consistent with active cohesin-mediated LE proceeding on both Y and U chromatin.

Overall, these observations highlight two different spatial configurations of Mmyco and Mpneumo exogenous chromosomes. On the one hand, the inactive Mmyco chromosome in G1 is compacted and positioned at the periphery of the nucleus, away from the active chromosome set. Upon S-phase and G2 arrest, the DNA is nevertheless compacted in a cohesin-dependent manner with no visible loops at discrete positions. On the other hand, the active Mpneumo chromosome behaves as endogenous yeast chromosomes, displaying discrete mitotic DNA loops in G2/M.

### Mosaic chimeric chromosomes display spontaneous, chromatin type-dependent DNA compartmentalization

The structural and functional features of the Y- and U-type chromatin are reminiscent of the euchromatin and heterochromatin compartments described along metazoan chromosomes ^24^. We wondered whether they can coexist on a single chromosome. Using CRISPR-Cas9, we first fused the Mmyco chromosome with the native yeast chromosome XVI (XVIfMmyco chromosome). We then induced translocations to generate two other strains with chromosomes harboring alternating U and Y chromatin regions as small as 50 kb (XVIfMmycot1 and XVIfMmycot2; **Fig. 4a**; **Extended Data** Fig. 6a-c Methods). ChIP-seq profiles of RNA PolII on Mmyco regions were similar before and after translocation, demonstrating that PolII binding is independent of the broader regional context (**Extended Data** Fig. 6d). The overall Hi-C contact maps of XVIfMmyco strain synchronized in G1 showed little differences compared to the parental Mmyco strain (but for the deleted and fused regions) (**Fig. 4b**, compare panel **i** with **ii**). The Y-type chr. XVI arm intermixes with the other 15 yeast chromosomes while the U-type Mmyco arm remains isolated from the rest of the genome, contacting subtelomeric regions (**Fig. 4b**). In the translocated strains, however, the alternating chromatin type regions resulted in striking checkerboard contact patterns within XVIfMmycot1 and XVIfMmycot2 chromosomes (**Fig. 4b**, panels **iii** and **iv**). This pattern reflects the fact that U-type regions make specific contacts over long distances, bypassing the Y-type regions found in between (**Fig. 4c**; **Extended Data** Fig. 6e G1). Similarly, Y-type regions of chromosome XVI also are also involved in specific Y-Y contacts over longer distances (**Fig. 4c**). Intra U and Y regions contact decay is also different in the two chromatin types, U chromatin being prone to longer range contacts in cis (**Extended Data** Fig. 6e). We then induced another translocation to integrate a segment of U chromatin in a *trans* position at the end of chromosome XIII and generated the corresponding Hi-C map (**Fig. 4d**). The correlation and 4C-like plot revealed an increase in inter-chromosomal contacts between Y chromatin segments with distant segments of itself, and in alternance with decreased contacts with Mmyco U chromatin segments (and vice versa), showing that the inactive U chromatin compartment can contain segments positioned on different chromosomes (**Fig. 4d**). These results unambiguously show that in yeast alternated or inter-chromosomal segments of active and inactive chromatin are spontaneously segregated within the nuclear space in G1 (**Fig. 4e, Extended Data** Fig. 6h), a phenomenon reminiscent of eu- and heterochromatin compartments found in higher eukaryotes ^24^.

**Figure 4.**
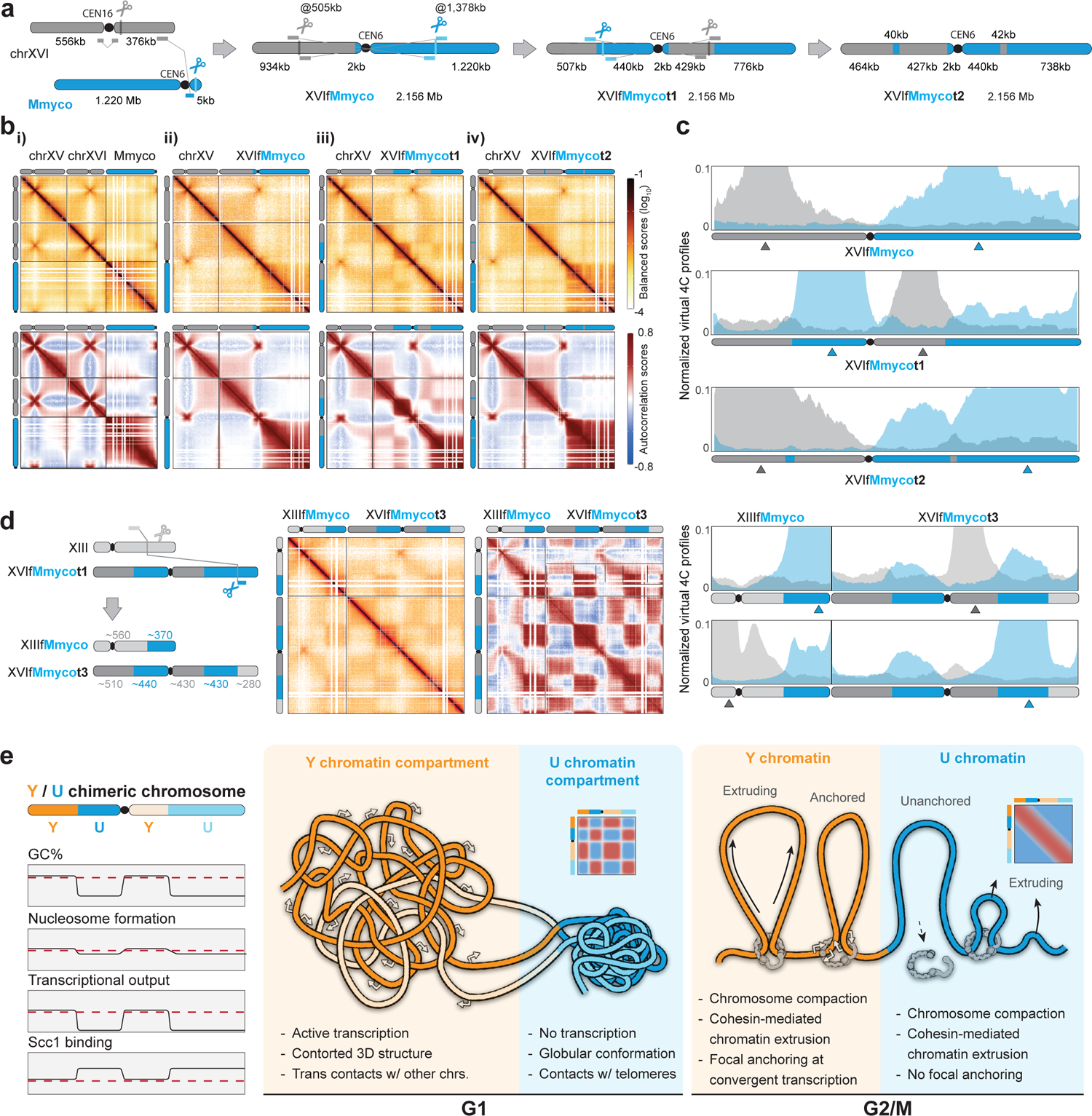
Compartmentalization of mosaic chromosomes composed of Y and U-type chromatin. **a,** Schematic representation of the CRISPR strategy used to generate the Mmyco mosaic chromosomes with alternating Y and U chromatin regions. **b**, Top: G1 Hi-C contact maps of chr. XV, XVI, and bacterial chromosomes in the Mmyco strain, and in translocated derivatives from panel **a**) (4kb resolution). Bottom: correlation matrices of the corresponding contact maps. **c,** Virtual 4C profiles of viewpoints (grey/blue arrows) located within yeast chr. XVI (in grey) or Mmyco chromosome segments (in blue), in XVIfMmyco (top), XVIfMmycot1 (middle) and XVIfMmycot2 (bottom) strains. **d,** Left: Schematic representation of the CRISPR strategy used to generate the Mmyco mosaic XVIfMmycot3 strain, with alternating yeast and Mmyco segments in chromosomes XIII and XVI. Middle: G1 Hi-C maps of mosaic chromosomes XIII and XVI in XVIfMmycot3 strain. The color scales for the normalized contact map and the correlation matrix are the same as in **b**. Right: Virtual 4C profiles of viewpoints located within yeast chr. XIII (in grey) or within yeast chr. XVI (in grey) or Mmyco chromosome (in blue) segments in XVIfMmycot3 strain. Grey/blue arrows:Y and U chromatin viewpoints position **e**, Schematic of a chimeric chromosome composed of alternating Y- and U-type chromatin. Left: composition of Y- and U-type chromatin. Middle: Spatial and functional organization in G1. Right: Organization of Y and U-type chromatin segments in G2/M.

In G2/M, the Scc1 deposition pattern, including centromere depletion, remained conserved along mosaic chromosomes (**Extended Data** Fig. 6d). Cohesin coating remained enriched over U-type Mmyco sequences, including over the 50 kb encompassed within 850 Mb of yeast DNA on XVIfMmycot2 chromosome. These results show that cohesin is loaded along inactive Mmyco regions independently of the proximity with the centromere (as along WT chromosomes) and independently of transcription, suggesting that the larger chromatin context plays little role in cohesin-mediated loading. In addition, Hi-C maps of the mosaic chromosomes in G2/M show that U-type compartments disappear upon cohesin-mediated compaction (**Extended Data** Fig. 6f). The analysis of distance-dependent contact frequency shows that all chromosome regions behave similarly at this stage (**Extended Data** Fig. 6e, G2/M). The only noticeable difference resulting from the fusion/translocation events in G2/M Hi-C contact maps consists in a DNA loop appearing over the boundary between Mmyco and the endogenous yeast sequence (**Extended Data** Fig. 6f,g). The two anchors of the loop are a strongly cohesin-enriched region already present on chr. XVI sequence, and another region slightly enriched in cohesin already present on the Mmyco sequence (**Extended Data** Fig. 6g, green arrows).

Together with observations from the previous section, these results show that Y and U chromatin types are both prone to dynamic cohesin-dependent DNA compaction and loop formation in G2/M phase (**Fig. 4e**) and further suggest that loop extrusion can proceed through the Y/U chromatin junction to form a loop bridging the nearest cohesin enrichment sites on each side ^35,36^. Therefore, as observed in multicellular organisms, LE-mediated metaphase chromosome compaction abolishes chromatin compartments to prepare chromosomes for segregation (**Fig. 4e)** ^33,37^.

## Discussion

### Chromatin composition and activity of bacterial DNA follows yeast sequence determinants

Several studies have shown that exogenous or random DNA segments in the yeast nucleus with a GC% relatively similar to that of yeast chromosome are actively transcribed ^38–41^. Here we explore the relationships between sequence composition and chromatin composition and activity by characterizing large, divergent bacterial DNA sequences in a eukaryotic cellular context into which they have not evolved. We find that *M. pneumoniae* and *M. mycoides* chromosomes integrated in yeast adopt one of two archetypes of chromatin that we called Y and U for yeast-like and unconventional. In both types, nucleosomes can assemble albeit with a longer NRL in the U chromatin (177 vs 160 bp), which also correlates with a low transcription. In G1, transcriptionally inactive U chromatin segments spontaneously collapse into a globule leading to the formation of a new nuclear compartment, whereas Y chromatin intermixes with yeast endogenous DNA. In G2/M, SMC cohesin complexes bind to both chromatin types and compact DNA through the extrusion of DNA loops ^6^. Along Y chromatin, these complexes accumulate at convergent transcription loci ^28,30^, whereas along U chromatin, the absence of active transcription leads to a homogeneous enrichment pattern.

The chromatin composition and activity experimentally measured over these exogenous chromosomes can be predicted using convolutional neural networks solely trained on yeast sequences, showing that exogenous DNA composition obeys a set of host rules based on sequence determinants that include GC content, dinucleotide composition but also complex sequence features. Sequence composition is therefore an important driver of the chromatin composition of exogenous sequences.

### Prokaryotic DNA contains some determinants of transcription orientation in eukaryote

Active transcription on bacterial chromosomes in yeast follows, on average, bacteria genes orientation (**Fig. 2**). Several mechanisms could account for this observation. Sequence determinants predating the divergence of eucaryotes and procaryotes, including GC and AT skews, are known to influence polymerase directionality ^47,48^. Alternatively, bacteria codon usage bias could result in enrichment in Nrd1-Nab3-Sen1 (NNS) binding sites along the ncRNA strand, leading to transcription termination ^39,49^. The Nonsense-Mediated Decay (NMD) complex, which destabilizes in the cytoplasm eukaryotic transcripts with long 3’ UTR, could also be more frequently activated by bacteria RNA transcribed in an antisense orientation. Eventually, the resulting conserved orientation could facilitate the domestication of exogenous sequences during horizontal transfer/introgression events between distant species. Whether these RNA molecules are translated, and peptides of bacterial origins exist in the yeast cell, remains to be determined. If so, they could provide a source of diversity and adaptation.

### Cohesin-mediated chromatin folding along non-transcribed DNA templates

It is proposed that cohesins expand DNA loops by an active process of loop extrusion until they encounter an obstacle and/or a release signal ^6,33^. For instance, the transcriptional repressor CTCF constitutes a cohesin roadblock to LE in mammals, but SMC-mediated loops have been identified in genomes lacking CTCF across domains of life. In yeast and other species, anchors are thought to be determined by a combination of convergent transcription, replication fork progression during S phase and/or the presence at these positions of stably bound cohesin promoting sister chromatid cohesion ^50–52^. The barely transcribed Mmyco chromosome is compacted by cohesin at G2/M, without discrete loop anchoring, suggesting that transcription is neither necessary for loading nor a primary driver for translocation. We propose that cohesins can move freely along this template without encountering significant obstacles, making it a suitable model for studying potentially blocking sequences or molecules.

### Compartmentalization of foreign DNA in host’s nucleus

We show that the introduction a foreign DNA with a lower GC content is sufficient to spontaneously promote the formation of a distinct inactive chromatin type, with increased NRL and a compact globular structure, all features reminiscent of mammalian heterochromatin. This chromatin forms a new compartment at the periphery of the budding yeast nucleus (**Fig. 5e**; Extended Data Fig. 6h), like the metazoan compartment “B” formed by inactive, H3K9me3/HP1-mediated heterochromatin, which is absent in yeast. The mechanisms behind compartmentalization remain unknown, but we hypothesize that the unmixing between U and Y chromatin could be due either to NRL change, protein composition and/or activity. Importantly, longer NRLs are found in heterochromatin in mammals, and are associated with histone H1 ^53^. In contrast, Y chromatin intermixes with active yeast DNA.

Depending on its sequence composition, a DNA molecule will therefore either compartmentalize or intermix with the host DNA. These different fates of the two chromatin types raise interesting evolutionary considerations. In the context of the invasion of the genome by exogenous mobile elements, this could lead either to the spontaneous isolation of inactive foreign DNA or, on the contrary, to the co-option of a new set of active sequences that could represent a reservoir of genetic innovations. This sequence-dependent mechanism may have contributed to the heterochromatinization of AT-rich transposable elements integrated in mammalian genomes.

## Authors Contributions

Conceptualization: CC, LM, JS, JM and RK. Methodology: LM, CC, JS, CL, JM, RK. Software: JS. Validation: CC, JS, LM. Investigation: LM, CC, with contributions from AP, AB, AgT MR and FB. Formal analysis: JS, AW, ER, CC, LM, FB, MR, AnT. Data Curation: JS, with contributions from CC, LM. Resources: GG, CL. Visualization: JS. Writing - original draft preparation: CC, JS, LM, JM and RK. Writing – Editing: all authors. Writing – Revisions: JS, LM, AW, JM, RK. Supervision: JM, RK. CC co-supervised a student. Funding acquisition: RK. Project Administration: RK.

## Acknowledgements

We are grateful to Bernard Dujon, Micheline Fromont-Racine, Alain Jacquier, Gianni Liti, Bertrand Llorente, Marcelo Nollmann, Cosmin Saveanu and Benoit le Tallec for fruitful comments on the work and the manuscript. This research was supported by the European Research Council under the Horizon 2020 Program (ERC grant agreement 771813) and Agence Nationale pour la Recherche (ANR-19-CE13-0027-02) to RK. RK, FB, JM and AnT also received support from Agence Nationale pour la Recherche (ANR-22-CE12-0013-01). CC was supported by a Pasteur-Roux-Cantarini fellowship. JS was supported by a postdoctoral ARC fellowship. We thank all our colleagues from the laboratory régulation spatiale des génomes for fruitful discussions. We especially thank Cyril Matthey-Doret for support during the earlier steps of the project. E. Turc and L. Lemée at Biomics Platform, C2RT, Institut Pasteur, Paris, France, are supported by France Génomique (ANR-10-INBS-09-09) and IBISA for processing and sequencing RNA samples.

