## Supplementary figures and methods for "Exogenous chromosomes reveal how sequence composition drives chromatin assembly, activity, folding and compartmentalization"

#### This file contains:

#### Supplementary results p. 2

#### Extended Data Figures 1 – 6 p. 3 - 12

#### Material and Methods p. 13 – 25

#### References p. 25 - 26

#### Supplementary results

**Sequence composition drives chromatin composition of exogenous chromosomes**

The different GC content values of Mmyco and Mpneumo prompted us to investigate if and how the sequence composition could explain the establishment of Y and U chromatin. To do so, we turned to convolution neural networks (CNN), a computational approach increasingly used in genomics to learn genome annotations from the underlying DNA sequence ^1,2^ (**Extended Data Fig. 3a)**. We trained CNNs to learn MNase-seq, and Scc1 or PolII ChIP-seq profiles using yeast chromosomes I to XV sequences (**Extended Data Fig. 3b,** Methods) ^3^. Inputs consist of 2-32kb sequences of the genome and the corresponding outputs consist of profile values at regularly spaced positions along the input sequence. For each strain, the trained CNNs predicted these three profiles with good accuracy on the held-out sequence of yeast chromosome XVI (correlation of 0.63, 0.82 and 0.68 for MNase, Scc1 ChIP and PolII ChIP signals respectively, computed from experimental and predicted average signals over non-overlapping genomic bins; **Fig. 2a-c,** see **Methods**), confirming that genomic sequence is a primary determinant of chromatin composition in yeast ^3–5^. Next, we applied the trained CNNs to Mmyco and Mpneumo sequences and found that the rules learned from yeast sequences were sufficient to accurately predict the features experimentally characterized on both exogenous sequences. Not only the predictions both qualitatively and quantitatively correctly recapitulate the features of Y chromatin on Mpneumo, but also the U chromatin features on Mmyco sequences, notably: (1) an increased NRL, (2) a broad Scc1 enrichment with no discrete peaks, and (3) a lower PolII coverage (**Fig. 2a-c,** **Extended Data Fig. 3c**).

To better characterize the link between GC content and chromatin composition and activity, we randomly generated artificial sequences of either 2 and 32 kb, with variable GC contents ranging from 0 to 100%, and predicted their average MNase-seq, Scc1 and PolII scores (**Fig. 2d**; **Methods**). All predicted signals display a strong variability with respect to the GC content. The MNase average signal was higher for intermediate GC % (i.e. between 30 and 50), whereas for higher and lower values it was strongly decreased. The cohesin signal showed a strong decrease for increasing GC %, and the PolII signal was predicted to be minimal stationary 20 and 40 % and increased for larger values. These predictions are in good agreement with experimental measurements derived from bacterial and yeast sequences cut into 100bp bins, suggesting that trained CNNs are indeed able to correctly predict average nucleosome occupancy, Scc1 and PolII over sequences with a wide range of GC content.

We then compared the dinucleotide enrichment on 100 bp genomic bins with experimental or predicted 10% highest coverage along chromosome XVI, Mpneumo or Mmyco, for MNase-seq, and Scc1 or PolII ChIP-seq. The same six dinucleotides (AG, AC, CT, CA, GT, TG, which are all formed by one strong and one weak nucleotide) are enriched in genomic loci with 10% strongest MNase signal, both in experimental and predicted coverages, across yeast and exogenous chromosomes (**Fig. 2e**), while the CpG dinucleotide is strongly depleted in these bins. This implies that this major determinant of nucleosome positioning in yeast can be used for the signal prediction in exogenous chromosomes. We also found that the Scc1 or PolII-specific experimental and predicted dinucleotide signatures mostly agree within each chromosome, albeit being weaker than the MNase-specific signature (**Extended Data Fig. 3d**), with the CpG dinucleotide strongly enriched in bins with high PolII coverage, suggesting that despite the fact that DNA methylation has disappeared in *S. cerevisiae*, yeast promoters are still enriched for CpGs.

Finally, we compared the efficiency of our CNN-based predictions of Mnase-seq, Scc1 and PolII ChIP-seq coverages with linear regression models using the GC% or the dinucleotide composition as predictor variables. We found that the predictive power of the CNN approach was greater than the linear models, indicating that other sequence features, more complex than GC% and dinucleotide composition, influence chromatin composition, and that these sequence determinants can be captured by convolution neural networks (**Fig. 2f**).

#### Replication and pairing of artificial chromosomes

#### DNA sequence composition and chromatin factors, including nucleosomes, are essential drivers of the replication timing, prompting us to investigate how this timing is established in Y and U chromatin ^6^. In yeast, replication initiates at the level of discrete, small autonomous replication sequences (ARS) positioned along all chromosomes. To investigate the replication timing profile of both bacterial chromosomes, which did not spontaneously evolve to contain these sequences, we used marker frequency analysis (MFA; Methods) ^7^. Whereas Mpneumo MFA profile exhibits DNA copy number variation indicative of the early or mid-S phase firing of multiple replication origins, MFA along Mmyco unveil the early firing of only the centromere-proximal ARS which was artificially integrated during chromosome assembly ^8^ (Extended Data Fig. 2a). The average copy number of the rest of the Mmyco chromosome appears flat and close to 1N in the MFA analysis. These observations confirm the role of chromatin composition in the replication profile as Y type chromatin found on the Mmyco chromosome is similar to the profile of endogenous chromosomes while the replication timing of U chromatin points at replication taking place late during S phase, most likely from cryptic ARS all along the AT-rich sequence ^9^. This pattern is reminiscent of the random and late-replication pattern displayed by the human inactive chromosome X ^10^.

#### DNA replication and SCC are closely related as cohesion is established during S phase through entrapment of sister DNA molecules by cohesin rings as the replication fork progresses ^11,12^. We therefore investigated SCC by performing image analysis of chromosome pairing (Methods; Extended Data Fig. 2b). Despite the strong enrichment in Scc1 deposition, SCC in Mmyco appears significantly reduced, in agreement with the flat replication pattern suggesting a late and random initiation of the replication process. On the other hand, the endogenous yeast chromosomes in Mmyco strain did not display a significant decrease in SCC, despite the apparent loss of cohesin at centromeres (Extended Data Fig. 3e).

#### Extended Data Figures and Legends

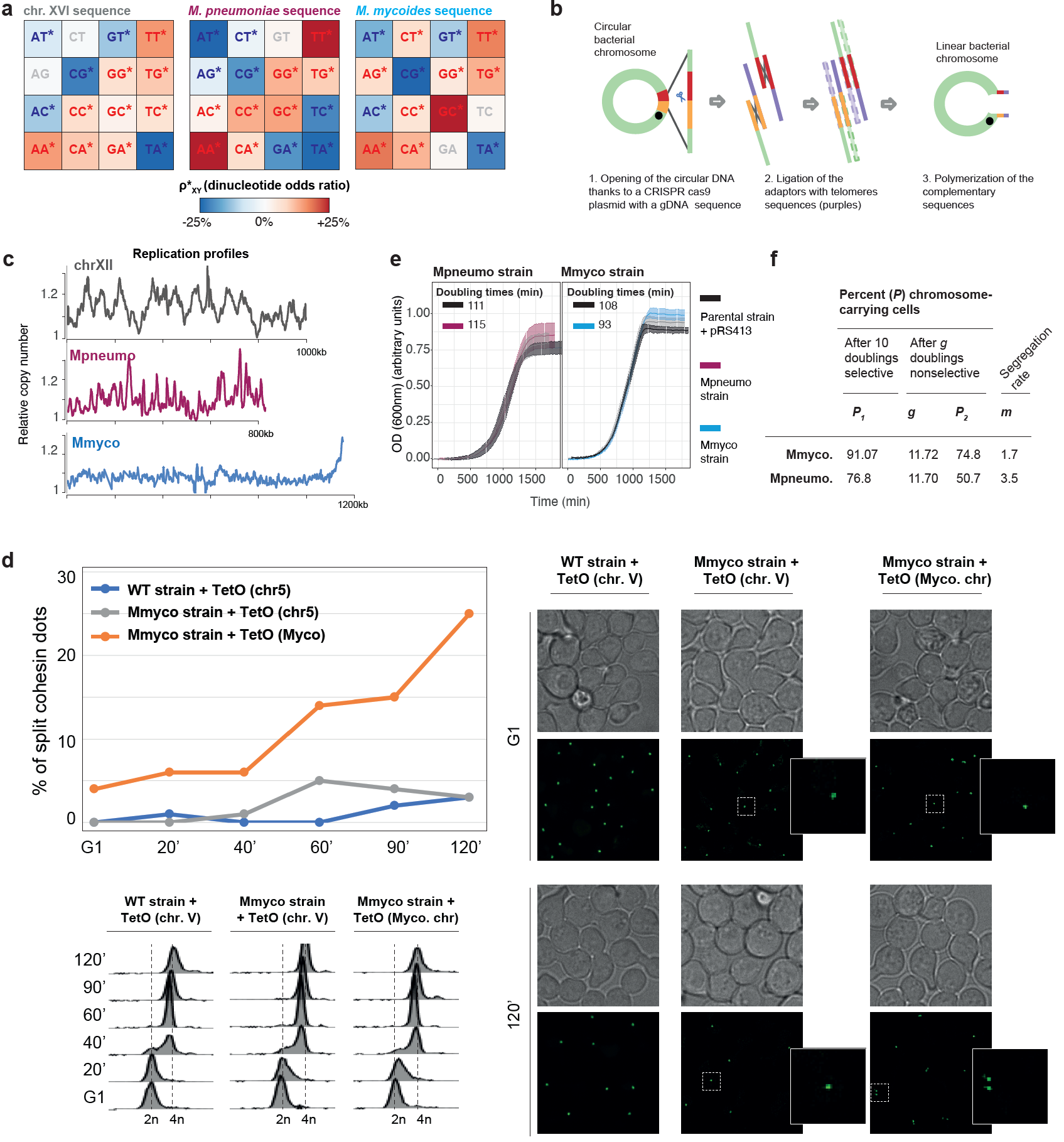

### **Extended Data Fig. 1. Characterization of Mpneumo and Mmyco strains metabolism.**

### **b,** Schematic representation of the CRISPR strategy applied to linearize and telomerize the circular bacterial genomes.

### **c,** MFA (replication profile) analysis of a representative WT yeast chromosome, and of Mmcyo and Mpneumo chromosomes.

**d,** Cohesin split dot assays in WT strain (TetO array in chr. V), or in Myco strain (TetO array in chr. V or Myco. chromosome). Top left: % of split cohesin dots in each strain upon G1-release; bottom left: cell ploidy upon G1-release; right: representative cohesin immuofluorescence imaging in each strain following G1-release.

**e,** Growth curves of WT, Mpneumo and Mmyco strains. For each strain, 3 independent cultures were performed. A pRS413 centromeric plasmid similar to the one onto which bacterial chromosomes were originally cloned is included in the WT strain as a control.

**f,** The chromosome stability and the segregation rate were measured as described in ^8^ (Methods). Yeast strains used are RSG_Y712 (Mmyco linear) and RSG_Y960 (Mpneumo linear). **P1**: % of bacterial chromosome-carrying cells in selective media. **g**: number of doubling. **P2**: of bacterial chromosome-carrying cells after 12 generations in non-selective media. **m**: segregation rate, i.e. % of plasmid-free segregants appearing in the final population after a single doubling.

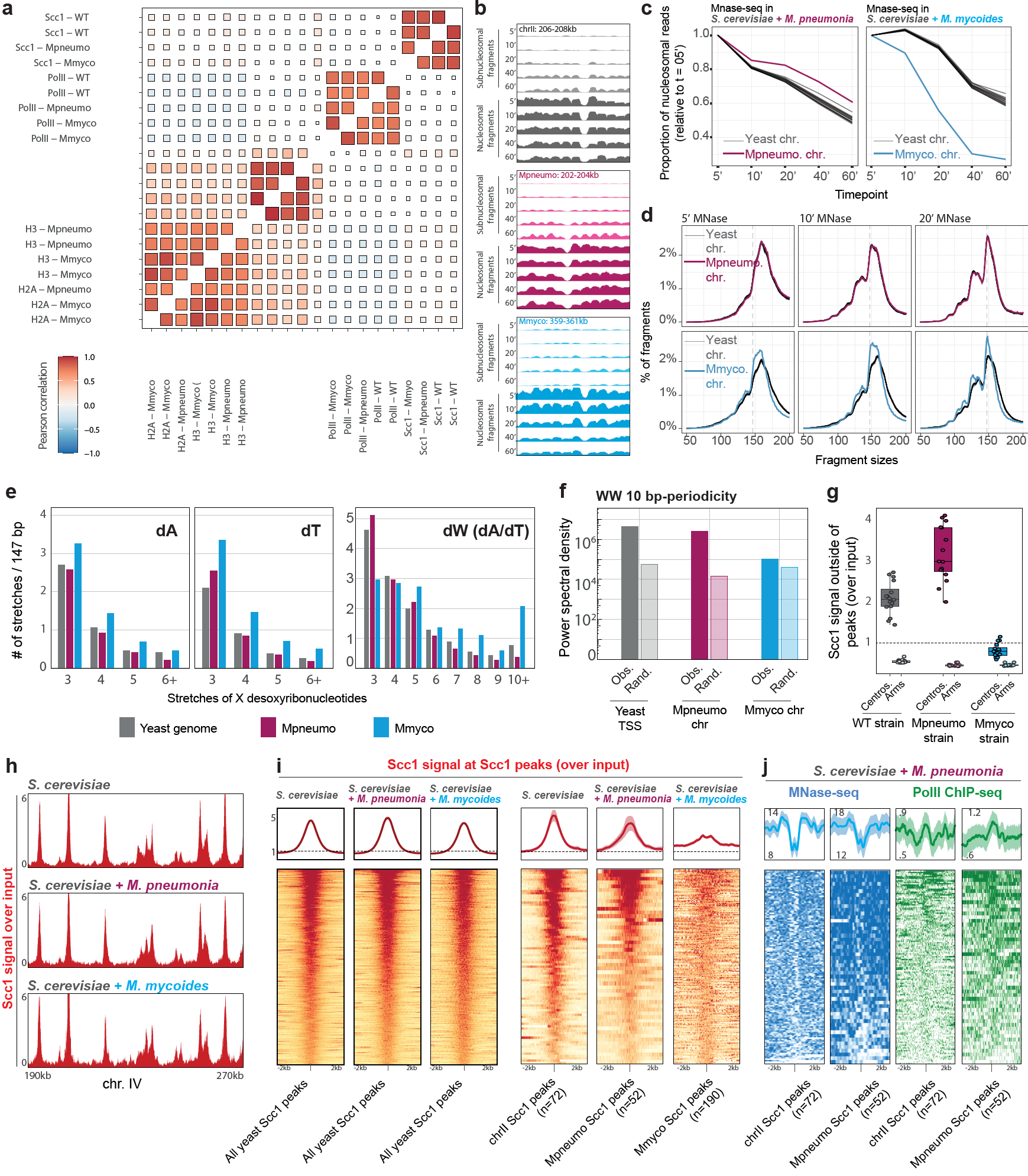

### **Extended Data Fig. 2. Replication of bacterial genomes in yeast.**

**a,** Pearson correlation between replicates of MNase and ChIP experiments. Bin size: 100bp.

**b,** Coverage of sub-nucleosomal (smaller than 130bp) or nucleosomal (between 130 and 165bp) fragments over 2kb-long genomic loci from chr. II, Mpneumo and Mmyco chromosomes, over an MNase digestion timecourse. All the tracks are displayed at the same scale (0-20 CPM).

**c,** Fraction of nucleosomal fragments versus sub-nucleosomal fragments in each chromosome at each timepoint, divided by the initial fraction at t = 5’.

**d,** Distribution of MNase-seq fragment sizes mapping over each chromosome after 5’, 10’ and 20’ of MNase digestion. The distribution over Mpneumo chromosome perfectly overlaps with the yeast chromosomes, whereas the distribution over Myco chromosome is shifted towards smaller fragments at each timepoint, indicating a more rapid degradation of nucleosomal fragments from Mmyco chromosome.

**e,** Number of poly-dA, poly-dT and poly-dW (dA/dT) stretches of various lengths in the yeast genome and over the Mpneumo and Mmyco chromosomes. Stretch numbers are scaled to 147bp.

**f,** Power spectral density (PSD) of WW 10-bp periodicity, in 300bp-long sequences centered over yeast TSSs or along the Mpneumo or Mmyco chromosomes. Random sequences were generated by shuffling actual sequences while preserving dinucleotide frequency.

**g,** Cohesin (Scc1) enrichment over yeast centromeres and yeast arms in WT, Mpneumo and Mmyco strains. Scc1 signal over arms was calculated outside of any Scc1 peak.

**h,** Representative 80kb window of Scc1 ChIP-seq deposition signal over yeast chr. IV in the WT, Mpneumo, and Mmyco strains.

**i,** Aggregated profile of Scc1 deposition centered at Scc1 peaks (+/- 2kb) called over all endogenous (left panels) or only chr. II (right panels) yeast chromosomes in the WT, Mpneumo and Mmyco strains.

### **j,** Aggregated profile of MNase-seq (blue) or PolII (green) deposition centered at Scc1 peaks (+/- 2kb) called over the yeast chromosome II (left) or the Mpneumo chromosome (right)

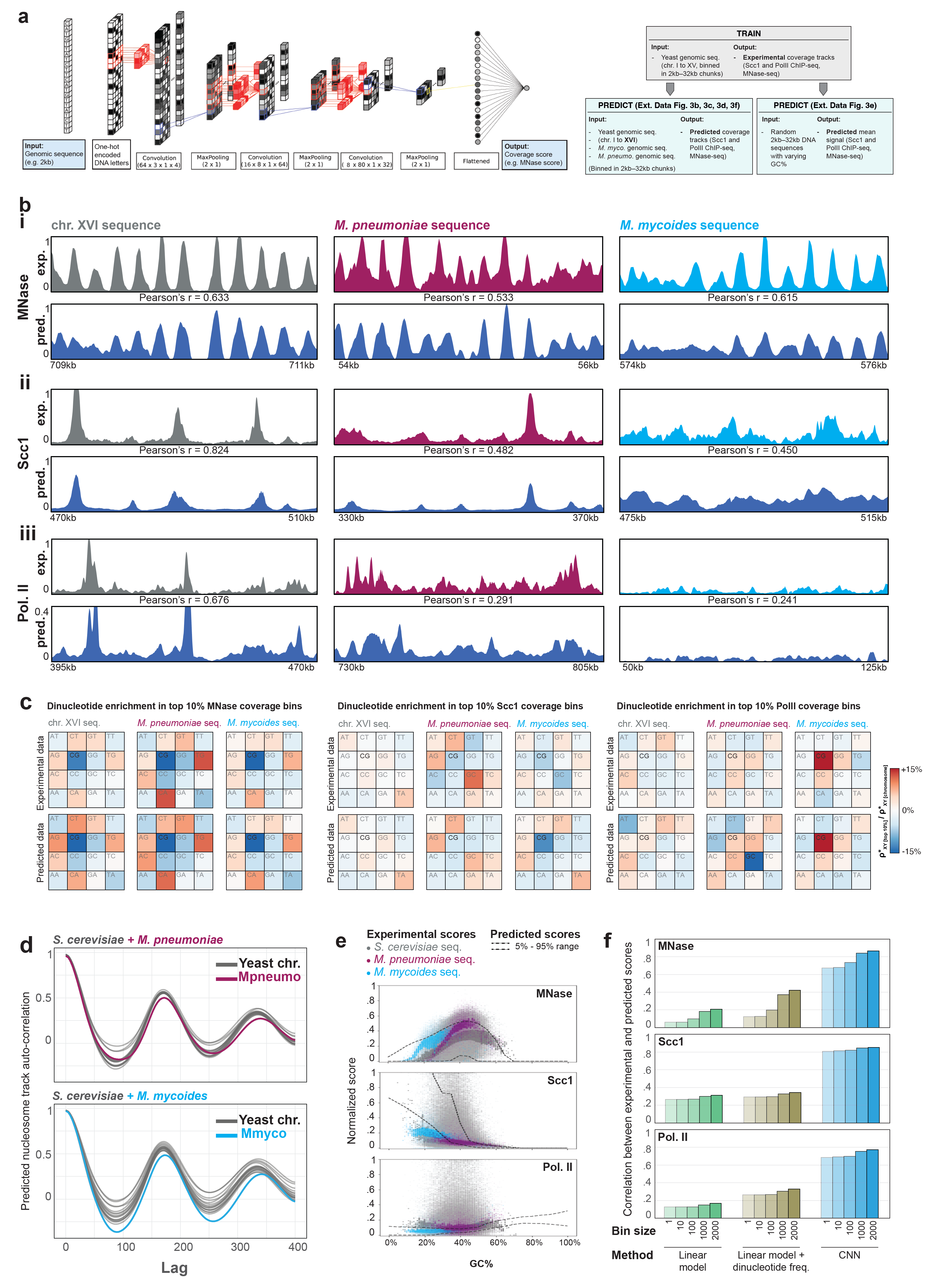

**Extended Data Fig. 3. Chromatin structuration of the bacterial chromosomes in yeast.**

**a, Left:** Schematic representation of the convolution neural network used in this study. The input of the network is a 2 to 30kb DNA sequence. The output is the corresponding value of the Mnase, Scc1 or PolII signal. Details about the size of the input/output and the architecture used are discussed in the Material and Methods. **Right:** Overall training/prediction strategy. We  train a CNN on sequences of  chr. I-XV to predict (i) genome-wide MNase-seq, Scc1 or PolII ChIP-seq coverage tracks over chrXVI and bacterial chromosomes, and (ii) the average MNase-seq, Scc1 or PolII ChIP-seq signal over 10kb random sequences with varying GC%.

**b,** Experimental (top) and NN prediction (bottom, dark blue) of nucleosome tracks or Scc1 and PolII ChIP-seq coverage tracks, over yeast chromosome XVI or over Mmyco and Mpneumo bacterial chromosomes in yeast. Signals have been smoothed using a sliding genomic window of 10 bp (MNase-seq) or 500 bp (ChIP-seq). For each chromosome, the Pearson correlation score was computed between experimental and predicted scores, using averaged scores over non-overlapping 10 bp bins for MNase or 500 bp bins for ChIP-seq. Bins with an average score lower than 0.01 were excluded.

**c,** Dinucleotide enrichment in genomic loci with 10% greatest MNase (left), Scc1 (middle) or PolII (right) ChIP-seq coverage (100 bp bins), extracted from experimental or predicted tracks over chromosome XVI, Mpneumo or Mmyco. The dinucleotide composition in these loci is compared to the chromosome-wide dinucleotide composition (ρ*(XY)_[chromosome]_ / ρ*(XY)_[chromosome]_).

**d,** Auto-correlation of the nucleosome tracks predicted over yeast + Mpneumo or yeast + Mmyco genome sequences. For each genome, the auto-correlation is computed for each chromosome separately (lag <= 400).

**e,** Dotted lines: 5%-95% range of predicted MNase-seq, Scc1 and PolII ChIP-seq scores in 2kb (MNase) or 30kb-long (ChIP-seq) random sequences with varying GC content. The experimental 100bp averaged scores measured along yeast, Mmyco and Mpneumo chromosomes are shown as colored dots (scores of the middle third for each GC% unit are in bold).

**f,** Correlation between experimental and predicted MNase-seq, Scc1 or PolII ChIP-seq data, using a linear model accounting for GC% only (green) or dinucleotide composition (kaki), or using CNNs (blue).

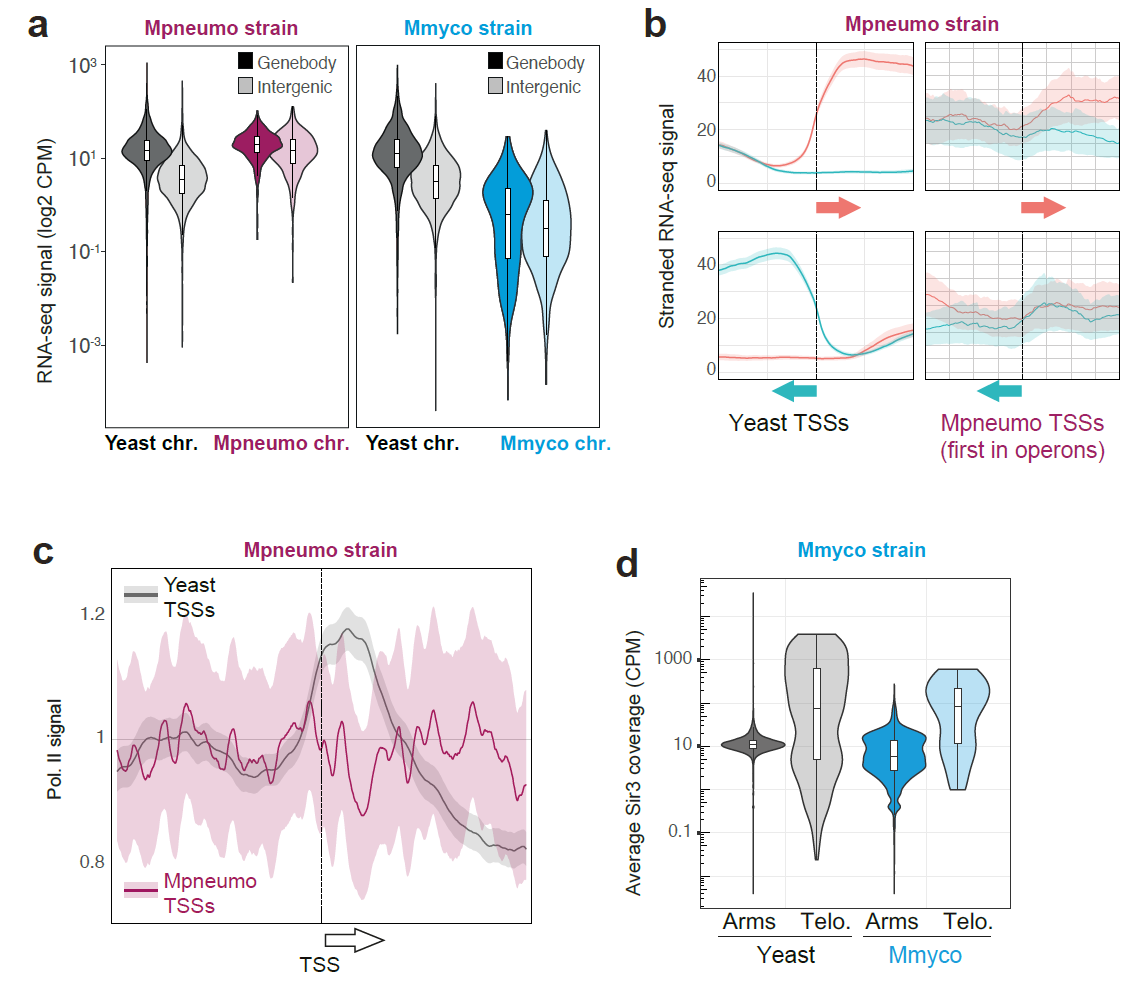

### **Extended Data Fig. 4. Transcription orientation of bacterial genomes in yeast.**

**a,** RNA-seq average signal over yeast or bacterial gene bodies and intergenic regions, in Mpneumo and Mmyco strains. Scores were normalized by the length of each genomic feature (CPM: counts per million of sequenced fragments).

**b,** Stranded analysis of RNA-seq data in Pneumo. strain. Pile-up of 1kb windows centered on transcription start sites (TSS) of genes either in the forward (Top) or reverse (Bottom) orientation. Left: endogenous yeast genes. Right: TSS of the first gene of annotated operons along the *M. pneumoniae* sequence.

**c,** PolII ChIP-seq coverage analysis in Pneumo strain. Pile-up of 2kb windows centered on transcription start sites (TSS). Grey: endogenous yeast genes. Purple: TSS of the first gene of annotated operons along the *M. pneumoniae* sequence.

**d,** Average Sir3 ChIP-seq signal over chromosome arms or telomeres (up to 3kb from end of chromosomes) in endogenous yeast chromosomes or in Myco. chromosome.

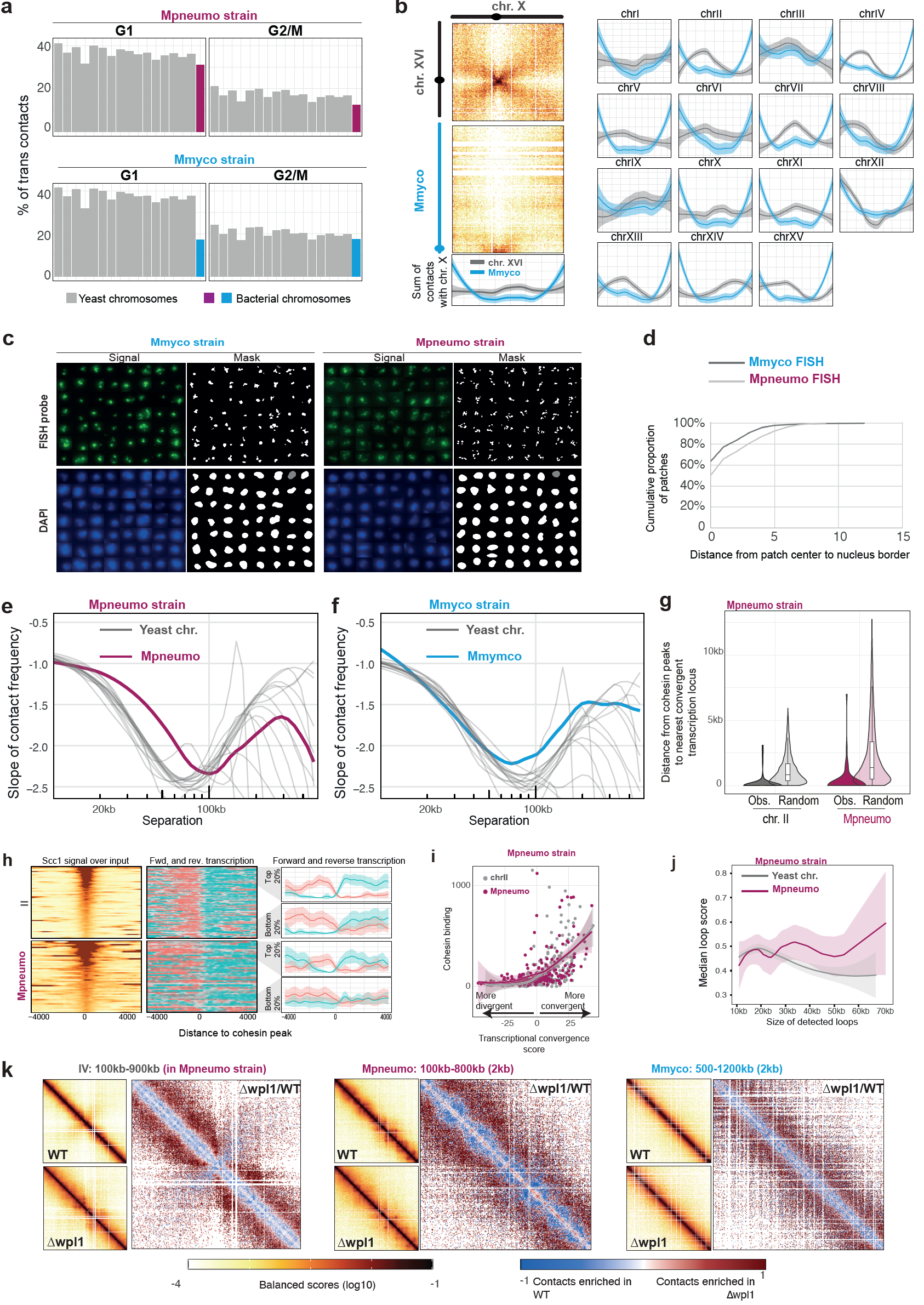

### **Extended Data Fig. 5. Folding of exogenic bacterial sequences within the yeast nucleus.**

**a,** % of total trans-contacts made by endogenous yeast chromosomes, Mmyco (top) or Mpneumo (bottom) bacterial chromosomes, in G1 or G2/M.

**b,** Left: quantification of contacts between the entire Mmyco chromosome (blue) and endogenous yeast chr. X (grey). Right: similar analysis for the other 15 endogenous yeast chromosomes. Note the increase of Myco. contacts at yeast telomeres.

**c,** Series of nuclei from Mpneumo or Mmyco fixed cells labeled with DAPI and hybridized with a fluorescent probe generated from either purified Mmyco (left) or Mpneumo (right) chromosome (top row: probe signal; bottom row: DAPI signal).

**d,** Distance between the patch of fluorescent signal from either the Mmyco or Mpneumo chromosomes and the nucleus border. Note that the Mmyco patches are located closer to the nucleus border than Mpneumo ones.

**e, f,** Slope of distance-dependent contact frequency of endogenous yeast chromosomes and bacterial chromosomes in G2/M Mpneumo (**c**) and Mmyco (**d**) strains.

**g,** Distance measured between cohesin peaks and their nearest convergent transcription locus, for peaks located in chr. II or in Mpneumo chromosome. Expected distances, measured after randomly shuffling the position of the cohesin peaks 100 times, are also shown.

**h,** Left: aggregated profile of Scc1 deposition centered at Scc1 peaks (+/- 4kb) called over chr. II (top) or Mpneumo (bottom), with peaks ordered by peak strength. Middle: corresponding stranded transcription tracks, colored according to their forward or reverse orientation. Right: for chr. II or Mpneumo chromosome, average forward and reverse transcription over the 20% strongest or 20% weakest Scc1 peaks.

**i,** Correlation between Scc1 (cohesin) binding and convergent transcription strength (see Methods) in chr. II and in Mpneumo chromosome.

**j,** Distance-dependent loop scores (computed using Chromosight ^13^) for loops along either endogenous (grey) or Mpneumo (purple) chromosomes.

**k,** For yeast chromosome IV, Mpneumo and Mmyco: Left, Hi-C contact maps of the endogenous yeast chromosome IV of the Mpneumo strain synchronized in G2/M in either WT and Wpl1 depleted cells (Δwpl1); Right, corresponding chr. IV ratio map (Δwpl1 over WT). Red (or blue) indicate enriched (or depleted) contacts in Δwpl1.

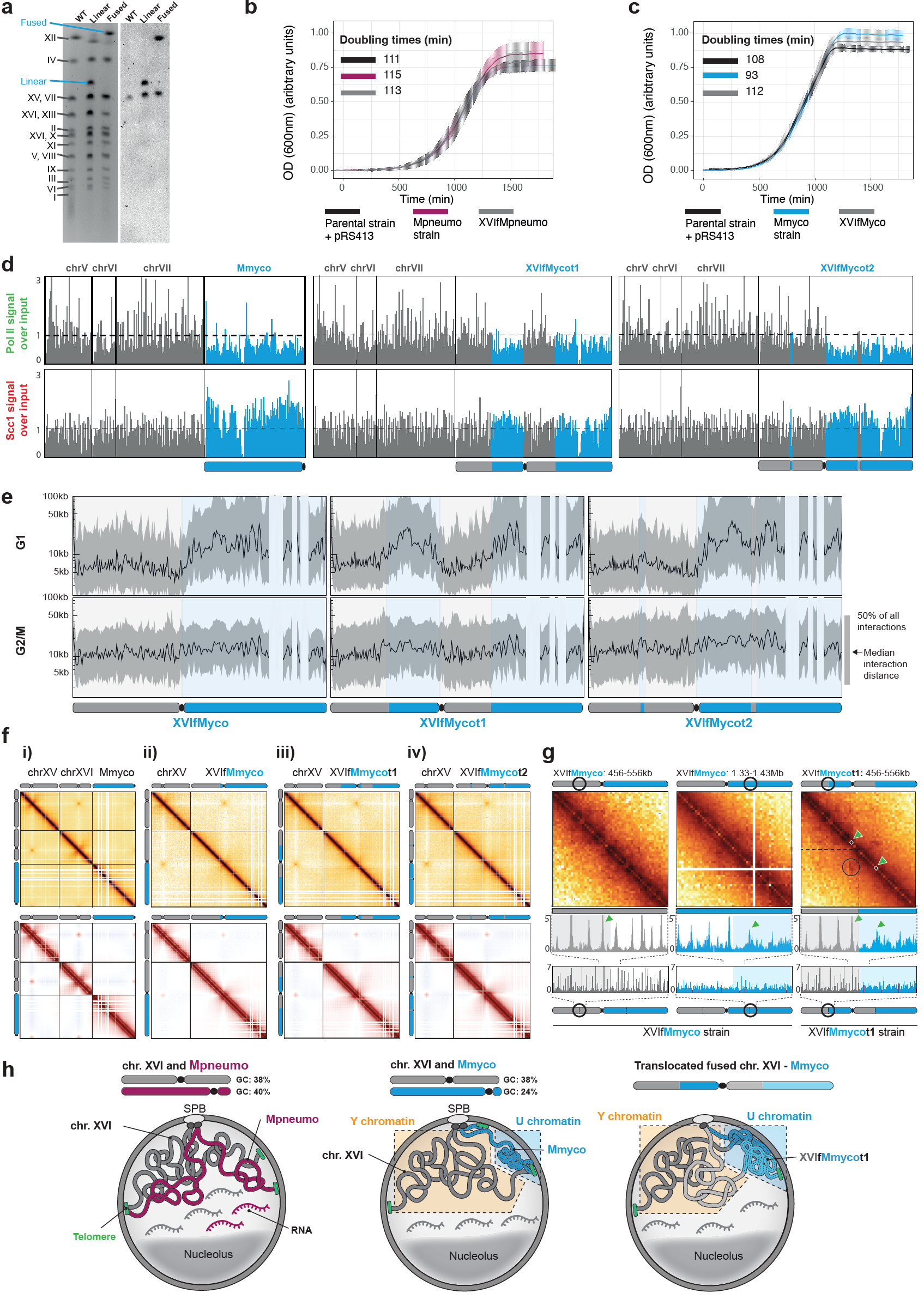

### **Extended Data Fig. 6. Compartmentalization of mosaic chromosomes composed of Y and U-type chromatin.**

### **a,** Left: pulsed-field gel electrophoresis of chromosomes from yeast strains containing the Mmyco chromosome, either linear or fused with endogenous yeast chr. XVI (XVIfMmyco strain). Right: Southern blot hybridization using a *his-3* probe present on the Mmyco chromosome sequence (note that *his-3* is also present on the endogenous chr. XV in the parental yeast strain).

**b,** Growth curves of WT, Mpneumo and XVIfMpneumo strains.

**c,** Growth curves of WT, Mmyco and XVIfMmyco strains.

**d,** PolII (top) and Scc1 (bottom) ChIP-seq deposition profiles along three representative yeast chromosomes and Mmyco chromosome (left) and mosaic chromosomes in XVIfMmycot1 (center) and XVIfMmycot2 (right) strains. Bin size: 10kb.

**e,** Average distance of interaction along the fused and mosaic Mmyco chromosomes: XVIfMmyco (left), XVIfMmycot1 (center) and XVIfMmycot2 (right). The shaded ribbon represents the interval between the 25% and 75% quantile of distance of interactions.

**f,** Top: G2/M Hi-C contact maps of chr. XV, XVI, and bacterial chromosomes in the Mmyco strain, and in its derivatives (i.e. the chr. XVI and Mmyco fusion, and the two strains with translocations resulting in alternating Y and U chromatin segments; Methods). Bottom: auto-correlation contact matrices in wt and mosaic chromosomes strains. The color scales are the same as in Fig. 5b.

**g,** Hi-C contact maps in G2/M of 100 kb window of either the XVIfMmyco (left and center) or XVIfMmycot1 (right) chromosomes, centered on the translocation position of the XVIfMmycot1 chromosome. In XVIfMmycot1, this window is effectively centered at the junction between the yeast chr. XVI segment (upstream) and the Myco chromosome segment (downstream). The Scc1 deposition profile measured by ChIP-seq in each strain is shown underneath each contact map. Green arrows: cohesin peaks flanking the junction between the yeast and the Myco segments in the XVIfMmycot1 chromosome.

**h,** Illustration of the behaviors and activity of Mpneumo (left panel) and Mmyco (middle and right) chromosome sequences integrated in the yeast genome. Note that yeast and Mpneumo chromosomes intermingle in a single nuclear compartment, whereas the Mmyco chromosome (independently or in a chimeric state within chr. XVI) is condensed and segregated at the nuclear periphery, thereby defining the U-type chromatin compartment.

#### Material and Methods

**Strains and medium culture conditions**

All yeast strains used in this study are derivatives of W303 or VL6-48N and are listed in the Table “Strain list” (**Table S1**). The original Mpneumo and Mmyco yeast strains carry the circular genomes of *Mycoplasma mycoides* subsp. *mycoides* strain PG1 ^14^ and *Mycoplasma pneumoniae* strain M129 ^8^, respectively. Yeast strains were grown overnight at 30°C in 150mL of suitable media to attain 4,2x10^8^ cells. Cells were grown in a synthetic complete medium deprived of histidine (SC -His) (0.67% yeast nitrogen base without amino acids) (Difco), supplemented with a mix of amino-acids, 2% glucose) or in in rich medium (YPD): 1% bacto peptone (Difco), 1% bacto yeast extract (Difco), and 2% glucose. Yeast cells were synchronized in G1 by adding α-factor (Proteogenix, WY-13) in the media every 30 min during 2h30 (1 µg/mL final). To arrest cells in metaphase, cells were washed twice in fresh YPD after G1 arrest and released in rich medium (YPD) containing Nocodazole (Sigma-Aldrich, M1404-10MG) during 1h30. Cell synchronization was verified by Flow-cytometry.

**CRISPR-Cas9 engineering**

We used a CRISPR–Cas9 strategy to linearize the circular bacterial chromosome present in the parental yeast strains. Note that the highly acrocentric structure of Mmyco corresponds to the only position that was cleaved among several tested to linearize the chromosome. We suspect that due to the high AT content the Crispr targeting remains relatively poorly efficient. Plasmids pML107 ^15^ or PAEF5 ^16^ carrying gRNA were co-transformed with 200bp DNA repair recombinant donor sequences carrying telomeric repeats seeds (Genscript Biotech Netherlands) (Figure Supp 1A).

We also applied a CrispR-Cas9 approach to concomitantly fuse chromosome XVI with the linear bacterial chromosomes and remove chromosome XVI centromere, as described in Luo et al. ^17^. The resulting chromosome (XVIfMmyco) as one bacterial DNA arm carrying U chromatin and one yeast DNA arm made of Y chromatin (**Fig. 6A**). Strain XVIfMmyco’s karyotype was verified using pulsed-field gel electrophoresis (**Fig. S6A**), and that it grows normally was verified by growth assay (**Fig. S6B, C**). Briefly, we used three gRNAs inserted in pGZ110 and pAEF5 ^16^ co-transformed with recombinant DNA donors sequences (Twist Biosciences). These donor sequences are designed so that upon recombination 1) centromere XVI is removed and 2) fusion takes place between subtelomeres of chromosome XVI and either Mmyco and Mpneumo right arms (**Fig. 6A**). The same strategy was applied to generate reciprocal translocations between chromosome XVI and Mmyco sequences of the XVIfMmyco chromosome (**Fig. 6A**). We generated alternating domains of Y and U chromatin along the XVIfMmyco chromosome using CRISPR-induced reciprocal translocations between the two arms, resulting in strains XVIfMmycot1 and t2 carrying alternating regions of U and Y type chromatin along a chromosome (**Fig. 6A**). The chromatin composition of these chimeric chromosome consists of [TEL - 500kb Y - 700kb U - CEN - 400kb Y - 500kb U - TEL] for XVIfMmycot1 and of [TEL - 500kb Y - 50kb U - 350kb Y - CEN - 650kb U - 50kb Y - 500kb U - TEL] for XVIfMmycot2.

gRNA sites were chosen to optimize Crispr targeting specificity using chopchop.cbu.uib.no ^18^ or CRISPOR ^19^. For all experiments, 100 ng of gRNA expression plasmids and ~400 ng of recombinant donors were co-transformed. After co-transformation, yeast cells were immediately plated on the corresponding selective media. Yeast cells were allowed to recover in fresh YPD for 2h before plating.

Linearization, fusion and translocations of chromosomes were verified by PCR, pulse-field gel electrophoresis and Hi-C.

**Liquid growth assays and segregational plasmid stability**

Growth rate of parental, Mmyco and Mpneumo strains carrying linear or fused chromosomes were assessed for independent clones in triplicates. Briefly, independent clones were grown overnight in selective medium (SC -HIS) to saturation at 30°C. The following morning the cultures were diluted to OD600 = 0.01 and inoculated in 96 well plates containing 100μl fresh SC -HIS. The cells were grown under agitation in a Tecan Sunrise plate reader at 30°C (Tecan). optical densities at 600 nm were recorded every 10 min to generate the growth curves for individual wells. The average of triplicates was plotted to compute the doubling time.

Segregational plasmid stability was measured as described in ^20^. Three individual transformants for each strain were inoculated into SC-HIS. P1 is the percentage of bacterial chromosome-carrying cells in selective media. It was determined from the ratio of viable colonies obtained by plating on YPD plates then replicated in selective medium (SC -HIS). To measure the chromosome stability, cultures were diluted into non-selective media and grown for X generations (g). After approximately 12 generations, the percentage of cells containing the bacterial chromosome (P2) was also determined from the ratio of viable colonies on rich medium (YPD) then replicated in selective medium. We then used P1, P2 and (g) number to calculate the segregation rate (m), which is defined as the % of plasmid-free segregants appearing in the final population after a single doubling.

**Pulsed-field Gel Electrophoresis and Southern blot hybridization**

Agarose plugs containing yeast chromosomes were prepared as described ^21^ and separated by clamped homogeneous electric field gel electrophoresis Rotaphor (Biometra) using the following parameters. Gel: 1% (SeaKemGTG); t=12 °C; buffer: 0.25 × Tris-Borate-EDTA; Program: [140 V, switch time: 300 s to 100 s, run time: 70 h].

DNA from the PFGE was transferred on a membrane. Southern blot was performed using digoxigenin-labeled DNA probes as described ^22^. We used the HIS3 gene (YOR202w) present near the Mmyco and Mpneumo centromere was used as target to validate the linearization and the fusion. Digoxigenin-labeled DNA probes were synthesized from the pRS413 plasmid with the following oligos: 3’CTACATAAGAACACCTTTGG5’ and 3’ATGACAGAGCAGAAAGCCCT5’.

**Hi-C procedure and sequencing**

Cell fixation with 3% formaldehyde (Sigma-Aldrich, Cat. F8775) was performed as described in Dauban et al. ^23^. Quenching of formaldehyde with 300 mM glycine was performed at 4°C for 20 min. Hi-C experiments were performed with a Hi-C kit (Arima Genomics) with a double DpnII + HinfI restriction digestion following manufacturer instructions. Samples were purified using AMPure XP beads (Beckman A63882), recovered in 120ul H2O and sonicated using Covaris (DNA 300bp) in Covaris microTUBE (Covaris, 520045). Biotinylated DNA was loaded on Dynabeads™ Streptavidin C1 (FISHER SCIENTIFIC, 10202333). Preparation of the samples for paired-end sequencing on an Illumina NextSeq500 (2x35 bp) was performed using Invitrogen TM Collibri TM PS DNA Library Prep Kit for Illumina and following manufacturer instructions.

**ChIP-seq**

The ChIP-seq protocol is described in ^24^. Experimental replicates (x3) were made for each condition. Briefly, cells of either *S. cerevisiae* or *Candida glabrata* were grown exponentially to OD600 = 0.5. 15 OD600 units of *S. cerevisiae* cells were mixed with 3 OD600 units of *C. glabrata* cells to a total volume of 45 mL for Scc1 calibration only. Cells were fixed using 4mL of fixative solution (50 mM Tris-HCl, pH 8.0; 100 mM NaCl; 0.5 mM EGTA; 1 mM EDTA; 30% (v/v) formaldehyde) for 30 min at room temperature (RT) with rotation. The fixative was quenched with 2mL of 2.5M glycine (RT, 5 min with rotation). The cells were then harvested by centrifugation at 3,500 rpm for 3 min and washed with ice-cold PBS. The cells were then resuspended in 300 mL of ChIP lysis buffer (50 mM HEPES KOH, pH 8.0; 140 mM NaCl; 1 mM EDTA; 1% (v/v) Triton X-100; 0.1% (w/v) sodium deoxycholate; 1 mM PMSF; 2X Complete protease inhibitor cocktail (Roche)) and transferred in 2mL tubes containing glass beads (ozyme, P000913-LYSK0-A.0) before mechanical cells lysis. The soluble fraction was isolated by centrifugation at 2,000 rpm for 3min then transferred to sonication tubes (Covaris milliTUBE 1ml, 520135) and samples were sonicated to produce sheared chromatin with a size range of 200-1,000bp using a Covaris sonicator. After sonication the samples were centrifuged at 13,200 rpm at 4°C for 20min and the supernatant was transferred into 700µL of ChIP lysis buffer. 80µL (27µl of each sample) of the supernatant was removed (termed ‘whole cell extract [WCE] sample’) and store at -80°C. 5ug of antibody was added to the remaining supernatant which is then incubated overnight at 4°C (wheel cold room). 50µL of protein G Dynabeads or protein A Dynabeads was then added and incubated at 4°C for 2h. Beads were washed 2 times with ChIP lysis buffer, 3 times with high salt ChIP lysis buffer (50mMHEPES-KOH, pH 8.0; 500 mM NaCl; 1 mM EDTA; 1% (v/v) Triton X-100; 0.1% (w/v) sodium deoxycholate;1 mM PMSF), 2 times with ChIP wash buffer (10 mM Tris-HCl, pH 8.0; 0.25MLiCl; 0.5% NP-40; 0.5% sodium deoxycholate; 1mM EDTA;1 mMPMSF) and 1 time with TE pH7.5. The immunoprecipitated chromatin was then eluted by incubation in 120µL TES buffer (50 mMTris-HCl, pH 8.0; 10 mM EDTA; 1% SDS) for 15min at 65°C and the supernatant is collected termed ‘IP sample’. The WCE samples were mixed with 40µL of TES3 buffer (50 mM Tris-HCl, pH 8.0; 10 mM EDTA; 3% SDS). All (IP and WCE) samples were de-cross-linked by incubation at 65°C overnight. RNA was degraded by incubation with 2µL RNase A (10 mg/mL) for 1h at 37°C. Proteins were removed by incubation with 10µL of proteinase K (18 mg/mL) for 2h at 65°C. DNA was purified by a phenol/Chloroform extraction. The triplicate IP samples were mixed in 1 tube and libraries for IP and WCE samples were prepared using Invitrogen TM Collibri TM PS DNA Library Prep Kit for Illumina and following manufacturer instructions. Paired-end sequencing on an Illumina NextSeq500 (2x35 bp) was performed. Libraries were performed in one or two biological replicates. When available, the duplicates were averaged for visualization. We calculated pairwise Pearson correlation scores between replicates and all showed high concordance.

**RNA-seq**

RNA was extracted using MN Nucleospin RNA kit and following manufacturer instructions. Directional mRNA library (rRNA removal) was prepared by the Biomics platform of Institut Pasteur, Paris and Paired-end sequencing (PE150) was performed by Novogene. Libraries were performed in three biological replicates and the triplicates were averaged for visualization. We calculated pairwise Pearson correlation scores between replicates and all showed high concordance.

**MNase-seq**

Each strain was grown to 10^7^ cells (OD600 of 0,8~) in 150 mL of SC -his medium at 30°C. Cells were fixed with 4 mL of formaldehyde 37% (1% final) for 20 minutes at room temperature. Cross-linking was stopped by adding 8 mL of 2.5M glycine (125mM final) for 30 minutes. The fixed cells were centrifuged, washed twice with cold phosphate-buffered saline (PBS 1X) and stored at -80°C. Once thawed, 900 µL of cells lysate obtained with a Precellys (Bertin Technologies) were recovered. 100 µL of 1X Micrococcal Nuclease Reaction Buffer (NEB, M0247S) and 5 µL of Bovine Serum Albumin (BSA, 20mg/mL) were added. The mix was divided into 10 tubes x 100 µL. 1 µL of Mnase enzyme (NEB, M0247S) at a concentration of 2,000,000 gel units/ml was added (2,000 units/sample final) in each tube. Samples were incubated at 37°C for varying times (0, 1, 2, 3, 5, 10, 15, 20, 40, and 60 minutes). 300 µL of Stop solution (10 µL of 0.5M EGTA pH 8.0, 30 µL of 20% SDS, 240 µL of H2O, and 20 µL Proteinase K 20mg/mL) was added to each tube. The tubes were then gently mixed and incubated at 65°C overnight. Samples were extracted with phenol/chloroform, and DNA was ethanol precipitated treated with DNase-free RNase. To evaluate the Mnase digestion kinetics, DNA samples were analyzed using an Agilent Tape Station. DNA was purified using 2.2X volume of AmpureXP beads and sequencing libraries prepared with Invitrogen TM Collibri TM PS DNA Library Prep Kit for Illumina following manufacturer instructions. Paired-end sequencing on an Illumina NextSeq2000 (2 x 50 bp) was performed**.**

**Replication MFA experiment**

Genomic DNA was prepared from asynchronous and G1 arrested cells in triplicates using Qiagen DNeasy Kit. Pellets were recovered, washed with cold 70% ethanol, air dried and dissolved in 50µl 1xTE. 100ng of soluble gDNA was transferred to sonication tubes and samples were sonicated to produce sheared chromatin with a size of about 300bp using a Covaris sonicator. Samples were purified using AMPure XP beads (Beckman A63882). Preparation of the samples for paired-end sequencing on an Illumina NextSeq500 (2x35 bp) was performed using Invitrogen TM Collibri TM PS DNA Library Prep Kit for Illumina and following manufacturer instructions.

**Processing of reads**

*Hi-C processing*

Reads were aligned and contact maps generated and processed using Hicstuff (https://github.com/koszullab/hicstuff). Briefly, pairs of reads were aligned iteratively and independently using Bowtie2 ^25^ in its most sensitive mode against their reference genome. Each uniquely mapped read was assigned to a restriction fragment. Quantification of pairwise contacts between restriction fragments was performed with default parameters: uncuts, loops and circularization events were filtered as described in ^26^. PCR duplicates (defined as multiple pairs of reads positioned at the exact same position) were discarded. Pairs were binned at 1kb resolution and multi-resolution balanced contact maps (in mcool format) were generated using cooler ^27^**.**

*MNase-seq processing*

For each timepoint, bowtie2 was used to align paired-end MNase-seq data on the appropriate genome reference. Only concordant pairs were retained, and fragments with a mapping quality lower than 10 were discarded. PCR duplicates were removed using samtools. Fragment coverage was normalized by library depth (CPM) and converted into a genomic track (bigwig) using deepTools.

*RNA-seq processing*

Bowtie2 was used to align paired-end RNA-seq data on the appropriate genome reference. Only fragments with an insert size shorter than 1kb were retained. Only concordant pairs were retained, and fragments with a mapping quality lower than 10 were discarded. PCR duplicates were removed using samtools. Fragment coverage was normalized by library depth (CPM) and converted into stranded or unstranded genomic tracks (bigwig) using deepTools.

*ChIP-seq processing*

For standard ChIP-seq, bowtie2 was used to align paired-end RNA-seq data on the appropriate genome reference. Only fragments with an insert size shorter than 1kb were retained. Only concordant pairs were retained, and fragments with a mapping quality lower than 10 were discarded. PCR duplicates were removed using samtools. When available, the input was similarly processed. Fragment coverage was normalized by library depth (CPM) and converted into a genomic track (bigwig) using deepTools. Input-normalized fragment coverage was also generated, when possible, using bamCompare from deepTools with “--scaleFactorsMethod readCount”. For histone ChIP-seq experiments, IP tracks were divided by input and log2-scaled. For calibrated Scc1 ChIP-seq, CPM-normalized fragment coverages were also multiplied by the ORi factor for calibration (WCE_glabrata_ x IP_cerevisiae_ / WCE_cerevisiae_ x IP_glabrata_, in which WCE_glabrata_ and IP_glabrata_ correspond to the number of paired reads that mapped uniquely on *C. glabrata* genome and same for *S. cerevisiae* reads). MACS2 2.2.7.1 was used to call peaks using the input alignment files as control.

**Analysis of genome-wide assays**

All downstream analysis steps were performed in R 4.1.2 / Bioconductor 3.16 using in-house scripts, unless mentioned otherwise.

*MNase-seq analysis*

To generate nucleosome tracks, “nucleosomal” fragments (between 130 and 165 bp) from the different MNase-seq timepoints were merged together then resized to a fixed 40bp length centered at the dyad. The resulting coverages were normalized to account for different numbers of nucleosomal fragments sequenced in the different MNase-seq timecourse experiments.

Nucleosome tracks were subsequently used to identify positioned nucleosomes as explained in ^28^. The linker DNA sequence was inferred as the distance between consecutive nucleosome dyads - 147bp. Auto-correlation was computed from the nucleosome tracks using the acf() function from the stats package in R, up to a maximum lag of 1000bp.

*ChIP-seq analysis*

Aggregated ChIP-seq profiles (average +/- 95% CI and heatmap) were plotted over windows centered at Scc1 peaks or TSSs after averaging coverage over 1bp-moving 200bp-wide rolling windows.

Median Scc1 coverage was calculated over 20 kb windows centered at yeast centromeres or over 300bp non-overlapping windows tiling the entire yeast chromosomes (excluding centromeres or Scc1 peaks).

Correlation scores between ChIP-seq replicates were calculated from the average coverage scores over 100bp tiled windows of yeast chromosomes.

*Sequence biases*

10-bp periodicity of dA, dT or dW dinucleotides in sequences up to 160 bp was computed over windows centered at yeast TSSs or tiling bacterial chromosome sequences, using “getPeriodicity'' function from periodicDNA ^29^, using ushuffle ^30^ to maintain a constant dinucleotide frequency in shuffled control sequences. K-mer occurrences were estimated over 147 bp windows centered at yeast TSSs or tiling bacterial chromosome sequences.

*Transcription convergence and Scc1 binding*

At every position *i* along the genome (every 10 bp), a *Dir_i_* directionality score was calculated as follows:

$${Dir}_{i} = \frac{\sum_{k = i}^{i + 200} ({RNA}_{fwd, k} - {RNA}_{rev, k})}{\sum_{k = i-200}^{i} ({RNA}_{fwd, k} - {RNA}_{rev, k})}$$

Genomic positions *j* for which ${Dir}_{j-101} > 0 \& {Dir}_{j-100} < 0$were then recovered and correspond to convergent or divergent transcription positions. For every transcription switch position *k*, a convergence score ${Conv}_{k}$ was subsequently computed as follows:

$${Conv}_{k} =\left[ \sum_{l = k - 2000}^{k} ({RNA}_{fwd,l} > 20) - \sum_{l = k - 2000}^{k} ({RNA}_{rev,l} > 20) + \sum_{l = k}^{k+2000} ({RNA}_{rev,l} > 20) - \sum_{l = k}^{k+2000} ({RNA}_{fwd,l} > 20) \right] \times100 / (4000 \times4)$$

Thus, ${Conv}_{k}$ scores range between -50 and 50 and a positive (negative) ${Conv}_{k}$ corresponds to a local genomic position $k$ of convergent (divergent) transcription. The relationship between ${Conv}$ convergence scores and average Scc1 coverage scores (over a 500bp window centered at the convergent position) was then computed.

*Replication MFA profile*

Replication profiles were analyzed using Repliscope version 1.1.1 as in ^9^*.*

*Hi-C contact maps*

All Hi-C contact and ratio maps were plotted with a log10 or linear scaling respectively, using “plotMatrix” function from HiContacts (<https://github.com/koszullab/HiContacts>).

*Loop detection*

Chromosight 1.3.1 was used to call loops de novo from contact maps binned at 1 kb and balanced with Cooler. Matrices were subsampled to contain the same total number of contacts. De novo loop calling was computed using the “detect” mode of Chromosight, with minimum loop length set at 2kb, percentage undetected set at 25 and pearson correlation threshold set at 0.315. Loop strength was quantified for each loop using the quantify mode of Chromosight and the mean loop score was calculated for each condition. Loop pile-ups of averaged 17kb windows were generated with Chromosight.

*Cis-trans ratio and P(s) analysis*

Cis-trans ratios were calculated using the “cisTransRatio” function from HiContacts. Contact probability as a function of genomic distance P(s) and its derivative were determined using pairs files (generated by hicstuff) and the “distanceLaw” function from HiContacts with default parameters, averaging the contact data by individual chromosomes.

*Virtual 4C profiles*

Virtual 4C profiles for 20kb-wide viewpoints were computed from 2kb-binned contact maps, using the “virtual4C” function from HiContacts. Contacts of entire chromosomes across the entire genome were manually computed using subsetting functions from HiContacts.

**Deep-Learning analysis**

*Models architectures and training*

Three different models were trained to learn either on the Mnase, PolII and Scc1 profiles from the underlying genomic sequence. The signals were pretreated as follows. First, we truncated the experimental profiles to a threshold corresponding to the 99th percentile of the profile distribution. We then divided all values in the truncated profile by the maximum value to get a signal between 0 and 1. For each position along the genome, a DNA sequence of length W was associated with a subset of n_out values from the corresponding profile to make the three datasets used for our deep learning framework. For each of these datasets, we used the yeast chromosomes I to XIII for training, IIV and XV for validation and XVI for test.

We implemented all the CNNs using the Keras library ^31^ and Tensorflow ^32^ as back-end. A RTX 2080 Ti GPU was used to improve the training speed. We used the adaptive moment estimation (ADAM) optimization method to compute adaptive learning rates for each parameter. The batch size was set to 512.

For the Mnase prediction task, our CNN architecture was similar to the one used in ^33^. It consists of three convolutional layers with respectively 64, 16 and 8 kernels of shape (3x1x4), (8x1x 64) and (80x1x16). A max pooling layer of size 2 and a ReLu activation function was applied after each of these three convolutions. Batch normalization and a dropout of 0.2 was applied after each convolution and the convolution stride was set to 1. Our model takes inputs of shape (2001, 1, 4), the last dimension representing the four nucleotides, and outputs a single value (i.e. n_out =1) corresponding to the value of the nucleosome profile in the middle of the input sequence.

For the PolII and Scc1 tasks, the architecture was modified as follows to take into account longer range influences ^34^. It consists of three convolutional layers all with kernels of shape (12x1x4), (8x1x64) and (80x1x16). Max Pooling layers of respective lengths of 8, 4 and 4 followed by ReLu activation were applied after each convolution. Four dilated convolution layers were then applied using 16 kernels of length 5 and dilatation values of respectively 2,4,8 and 16. For PolII, the input sequence length was set to 2048 bp and the output length n_out to 16, corresponding to the 16 values of the signal found every 128 bp over the 2,048 bp input sequence. For Scc1, the input sequence length was set to 32768 bp and the output length n_out was 256, corresponding to the 256 values of the signal found every 128bp over the 32,768 bp input sequence.

The loss function used is the sum of the Pearson's dissimilarity (1-correlation) between the prediction x and the target y and the mean absolute error (MAE) between them (loss = MAE(x, y) + 1 - corr(x, y)). This loss function has been previously shown to enable both a faster and a more accurate convergence ^33^.

An early stopping procedure was applied during training to prevent models from overfitting. The loss function was calculated on the validation set at every epoch to evaluate the generalisability of the model. The training procedure was stopped if the validation loss did not decrease at all for 6 epochs and the model parameters were set back to their best performing value. The training procedure usually lasted for 15 to 20 epochs.

*Predictions*

Predictions were done on all chromosomes, including the chromosomes from the training, validation and test sets as well as exogenous bacterial chromosomes.

In order to characterize the influence of the GCc content of the sequences on the overall profile heights, we generated 10000 random 2kb (MNase, PolII) or 32kb (Scc1) sequences and predicted each of the three profiles on these sequences.

*Comparison with experimental data*

For each chromosome, the Pearson correlation score was calculated between experimental measurements and predictions, using averaged scores over non-overlapping 10 bp bins for MNase-seq or 500 bp bins for ChIP-seq. Bins with an average experimental or predicted score lower than 0.01 were excluded from the correlation analysis.

**Imaging and analysis**

*Two-dots assay (SCC)*

Strains yLD126-36c, FB176 and FB200 were inoculated and grown overnight in SC-MET medium. The next day cultures were diluted to OD600=0.2 in SC-MET. After 3h of exponential growth, alpha factor (10 ul at 5mg/ml) was added every 30 min for 2 h. G1-arrested cells were released into YPD (plus 2 mM methionine), and aliquots sampled for FACS and imaging analysis.

*FISH experiments*

FISH experiments were performed as described in Gotta et al.^35^, with some modifications ^28^. The probes were obtained by direct labeling of the bacterial DNA (1.5 µg) using the Nick Translation kit from Jena Bioscience (Atto488 NT Labeling Kit), the labeling reaction was performed at 15°C for 90 min. The labeled DNA was purified using the Qiaquick PCR purification kit from Qiagen, eluted in 30 µl of water. The purified probe was then diluted in the probe mix buffer (50% formamide, 10% dextran sulfate, 2× SSC final). 20 OD of cells (1 OD corresponding to 107 cells) were grown to mid–logarithmic phase (1–2 × 107 cells/ml) and harvested at 1,200 g for 5 min at RT. Cells were fixed in 20 ml of 4% paraformaldehyde for 20 min at RT, washed twice in water, and resuspended in 2 ml of 0.1 M EDTA-KOH pH 8.0, 10 mM DTT for 10 min at 30°C with gentle agitation. Cells were then collected at 800 g, and the pellet was carefully resuspended in 2 ml YPD - 1.2 M sorbitol. Next, cells were spheroplasted at 30°C for 10 minutes with Zymolyase (60 µg/ml Zymolyase-100T to 1 ml YPD-sorbitol cell suspension). Spheroplasting was stopped by the addition of 40 ml YPD - 1.2 M sorbitol. Cells were washed twice in YPD - 1.2 M sorbitol, and the pellet was resuspended in 1 ml YPD. Cells were put on diagnostic microscope slides and superficially air dried for 2 min. The slides were plunged in methanol at -20°C for 6 min, transferred to acetone at -20°C for 30 s, and air dried for 3 min. After an overnight incubation at RT in 4× SSC, 0.1% Tween, and 20 μg/ml RNase, the slides were washed in H20 and dehydrated in ethanol 70%, 80%, 90%, and 100% consecutively at -20°C for 1 min in each bath. Slides were air dried, and a solution of 2× SSC and 70% formamide was added for 5 min at 72°C. After a second step of dehydration, the denatured probe was added to the slides for 10 min at 72°C followed by a 37°C incubation for 24h in a humid chamber. The slides were then washed twice in 0.05× SSC at 40°C for 5 min and incubated twice in BT buffer (0.15 M NaHCO3, 0.1% Tween, 0.05% BSA) for 30 min at 37°C. For the DAPI staining, the slides were incubated in a DAPI solution (1µg/ml in 1× PBS) for 5 minutes and then washed twice in 1× PBS without DAPI.

*FISH image analysis*

Images were acquired on a wide-field microscopy system based on an inverted microscope (Nikon TE2000) equipped with a 100×/1.4 NA immersion oil objective, a C-mos camera and a Spectra X light engine lamp for illumination (Lumencor, Inc). The system is driven by the MetaMorph software (Molecular Devices). The axial (z) step is 200 nm and images shown are maximum intensity projection of z-stack images (MIP). Quantifications were done on the MIP images using ilastik for segmentation and Fiji for analyses of the particles.

**Data availability**

Sample description and raw sequences for all figures are accessible on GEO database through the following accession number: GSE217022. Go to:

<https://urldefense.com/v3/__https://www.ncbi.nlm.nih.gov/geo/query/acc.cgi?acc=GSE217022__;!!JFdNOqOXpB6UZW0!s5OExaPiqjyc6NPF5fsCYTc8f3K4hsS_-t2_EXDnJ3RRHRukokAjjRoaBjocstAzCXasmp3PSgugmpjXb4UDP3FQuw$>

Enter token *arepcamenvafhuz* into the box.

Source data are provided with this paper.

**Code availability**

All custom-made code of the analysis of sequencing data is available online [https://github.com/koszullab/](https://github.com/koszullab/T7_promoter_analysis)

Open-access versions of the programs and pipeline used (Hicstuff) are available online on the github account of the Koszul lab Hicstuff ([https:/](https://github.com/koszullab/T7_promoter_analysis)/[github.com/koszullab/hicstuff](http://www.github.com/koszullab/hicstuff)) version 3.1.2, Chromosight (version 1.4.1 available online at https://github.com/koszullab/chromosight/), Bowtie2 (version 2.4.5 available online at http://bowtie-bio.sourceforge.net/bowtie2/), SAMtools (version 1.9 available online at http://www.htslib.org/), Bedtools86 (version 2.29.1 available online at https://bedtools.readthedocs.io/en/latest/content/installation.html) and Cooler (versions 0.8.7–0.8.11 available online at <https://cooler.readthedocs.io/en/latest/>).

All deep learning codes are available at:

<https://gitlab.in2p3.fr/mnhn-tools/npc-profile-prediction-on-exogeneous-genomes/-/tree/master/models>

Any additional information, including custom-made code required to reanalyze the data reported in this paper, is available from the lead contact upon request.
